# A new fMRI quality metric using multi-echo information: Theory, validation and implications

**DOI:** 10.64898/2026.03.19.712948

**Authors:** Javier Gonzalez-Castillo, César Caballero-Gaudes, Daniel A. Handwerker, Peter A Bandettini

## Abstract

Consistent, high-quality data is key to the success of fMRI studies given the many confounding factors and undesired signals that contaminate these data. Several quality assurance (QA) metrics exist for fMRI (e.g., temporal signal-to-noise ratio (TSNR), percent ghosting, motion estimates), but none of them leverage relationships between echoes that are part of multi-echo (ME) fMRI acquisitions. Here, we fill this gap by proposing a new QA metric for for ME-fMRI that quantifies the likelihood a given ME scan is dominated by BOLD (Blood Oxygenation Level-Dependent) fluctuations. We refer to this metric as *p_BOLD_*; the probability of the signal change being primarily BOLD contrast-dominated. Having an estimate of overall BOLD weighting – both before and after preprocessing - is meaningful because BOLD is the intrinsic contrast mechanism used in fMRI to infer neural activity. We introduce *p_BOLD_* to the neuroimaging community by first describing the theoretical principles supporting the metric. Next, we validate *p_BOLD_* efficacy using a small dataset (N=7 scans) of constant- and cardiac-gated scans that have distinct levels of contributing BOLD fluctuations. Third, we apply *p_BOLD_* to a larger publicly available ME dataset (N=439 scans), to evaluate six different pre-processing pipelines, and show how *p_BOLD_* provides complementary information to TSNR. Our results show that ME-based denoising increases both *p_BOLD_* and TSNR relative to basic denoising; however, including the global signal (GS) as a regressor only improves TSNR, but worsens *p_BOLD_*. Further analyses looking at the BOLD-like characteristics of the GS and its relationship to cardiac and respiratory traces suggest that the observed decrease in *p_BOLD_* is likely due to a decrease in BOLD fluctuations of neural origin contributing to the GS, and not due to contributions from other physiological BOLD fluctuations (i.e., respiratory and cardiac function). Finally, we also demonstrate how *p_BOLD_* can be applied as a data quality metric, by showing how higher *p_BOLD_* results in better ability to predict phenotypes based on whole-brain functional connectivity matrices.

## Introduction

Data quality strongly determines the success of any fMRI study (Reynolds et al., 2023). For example, unmodeled noise hinders our ability to detect small, yet meaningful, effects (Gonzalez-Castillo et al., 2015, 2012). Physiological confounds artifactually inflate functional connectivity estimates (Korponay et al., 2024). Systematic differences in head motion across populations result in incorrect inferences regarding brain development (Power et al., 2012); and it can also affect clinical conclusions (Makowski et al., 2019). Hardware instabilities can be particularly detrimental for resting-state data (Jo et al., 2010). Left-right orientation errors in data headers, reported in several large publicly available datasets (Glen et al., 2020), have consequences for functional laterality. Finally, inadvertent inconsistencies in acquisition parameters (e.g., repetition time, spatial resolution, spatial coverage, flip angle, etc.) can have negative interactions with pre-processing (e.g., incorrect TR voids temporal filtering) and explain sensitivity issues (e.g., echo time determines BOLD weighting in the data) for affected scans. While several pre-processing steps (e.g., motion correction, nuisance regression, smoothing) are designed to capture and correct these issues, they are not always possible. Incorporating comprehensive quality assurance protocols into everyday research practices is as important as working with clear research hypotheses or ensuring the soundness of statistical methods.

Over the years, and specially with the advent of large publicly available datasets, the fMRI community has become increasingly proactive at facilitating QA protocols for fMRI data. Early efforts focused on the definition of summary metrics that report issues associated with excessive head motion (Cox and Jesmanowicz, 1999; Oakes et al., 2005), drifting artifacts (Friedman and Glover, 2006), ghosting artifacts (Reeder et al., 1997), or compromised sensitivity (Bandettini et al., 1993); among others. More recently, mainstream functional neuroimaging analysis packages have streamlined the generation of comprehensive QA reports to facilitate effective identification of quality issues in large datasets. Examples of this include AFNI QA reports (Reynolds et al., 2023), CONN QA reports (Morfini et al., 2023), and NiPrep’s MRIQC reports (Provins et al., 2023); among others. One aspect of QA protocols that remains underdeveloped is that of QA metrics purposely tailored for multi-echo (ME) fMRI data—that is, metrics that leverage ME signal models to provide data quality insights not captured by traditional QC measures such as tSNR.

Multi-echo fMRI protocols concurrently acquire two or more timeseries per voxel; each at a different echo time (TE): the temporal offset between excitation pulses and the center of k-space readout windows. ME acquisitions were proposed during the early days of fMRI to experimentally explore BOLD weighings (Gati et al., 1997; Menon et al., 1993) and to improve sensitivity in dropout regions. Yet, due to prohibitive tradeoffs in acquisition time (i.e., acquiring more echoes resulted in longer repetition times, especially when collecting many slices), the general unavailability of turnkey ME sequences, and the uncertainty revolving how to best process multi-echo EPI time series, most researchers chose to continue with single echo acquisition. In the past 15 years, ME has begun to gain new momentum as an insightful and straightforward processing approach was introduced (Kundu et al., 2011). Other factors related to the recent resurgence of ME include the improved availability of ME/multi-band sequences at many research centers, the creation of the tedana denoising package (DuPre et al., 2021), and recent reports demonstrating its advantages over single-echo data for static (Lynch et al., 2020) and dynamic functional connectivity analysis (Cohen et al., 2021), deep phenotyping (Lynch et al., 2021), tasks involving overt recall inside the scanner (Gilmore et al., 2022),tasks highly contaminated with task-correlated noise (e.g., breath-hold (Moia et al., 2021), hand-grasping (Reddy et al., 2024)) and when performing hemodynamic deconvolution (Caballero-Gaudes et al., 2019; Uruñuela et al., 2024).

Despite this increase in ME adoption, no work to date has focused on the establishment of easily interpretable QA metrics unique to this type of data. While summary figures of the spatiotemporal variability of the fits for T_2_^*^ and S_o_ decay models are included in the tedana reports, these only provide a qualitative overview of one aspect of the data (i.e., the quality of the decay model fits) and are not amenable to efficient assessment of data quality in large ME datasets. To fill this gap, here we propose a novel QA metric specific to ME data called *p_BOLD_*. This metric was defined with the following principles in mind:

1. <u>Clear physical interpretation</u>: *p_BOLD_* quantifies the likelihood of a given scan being dominated by *T*_*2*_^*\**^ fluctuations (including BOLD), as opposed to net magnetization (S_o_) fluctuations. Having data dominated by BOLD fluctuations is desirable as BOLD is the primary intrinsic contrast mechanism used in fMRI to infer the location and timing of neural activity. That said, it is worth noting that not all BOLD fluctuations are expected to indicate neural activity. BOLD fluctuations can also be caused by variations in physiological processes (i.e., respiratory and cardiac functions) that also affect blood oxygenation levels (Krüger and Glover, 2001a; Triantafyllou et al., 2011) . As such, *p_BOLD_* ought to always be interpreted in the context of potential contributions from these other physiological processes. We consider this at length in the discussion section.
2. <u>Properties of a probability value</u>: by definition, *p_BOLD_* can only take values between zero and one. Values close to one indicate a high probability of the data being 100% BOLD (or BOLD dominated), values close to zero indicate otherwise.
3. <u>Based on TE dependence profile</u>: *p_BOLD_* relies on BOLD and *S*_*o*_ fluctuations having well differentiated behaviors across TEs: the amplitude of BOLD fluctuations is dependent on TE, while that of S_o_ fluctuations is not. This is the same principle at the core of echo-based denoising techniques such as ME-ICA (Kundu et al., 2014) and tedana (DuPre et al., 2021), and hemodynamic deconvolution methods tailored for ME-fMRI like ME-PFM (ME-Paradigm Free Mapping; (Caballero-Gaudes et al., 2019)).

The rest of this manuscript is organized as follows: First, in the theory section we start by showing how an ME signal model based on MRI signal equations help us differentiate BOLD and S_o_ fluctuations. Then, we describe how that model can be exploited to quantify the likelihood of a given scan being dominated by BOLD fluctuations. For this, we rely on inter-regional covariance profile across different echoes. We finalize the theory section with a step-by-step description of how to compute *p_BOLD_*.

Next, to confirm that the proposed theoretical framework behaves as expected on real fMRI data, we compute *p_BOLD_* on resting-state scans from a small sample of 7 subjects for whom we collected data in two ways: 1) using a constant repetition time (TR, constant-gated scans); and 2) using a non-constant repetition time linked to the peak of the cardiac cycle (cardiac-gated scans). Un-preprocessed cardiac gated data are expected to be S_o_ dominated given that irregular TRs introduce strong net magnetization effects on the signal. Conversely, low motion, tedana denoised regularly-sampled data is expected to be dominated by BOLD effects. This empirical evaluation confirms our expectations: *p_BOLD_* is significantly lower for cardiac-gated data as compared to low-motion, fully pre-processed constant-gated data.

We move next to real application scenarios where we test the efficacy of *p_BOLD_*, relative to that of TSNR, for selecting an optimal pre-processing pipeline, identifying individual scans with quality issues, and forecasting the strength of brain-wise associations. We use a publicly available large-N dataset (Spreng et al., 2022). We call this second dataset the “evaluation” dataset throughout the rest of the manuscript. Using 439 scans (from 221 subjects) in the evaluation dataset, we demonstrate that tedana denoising produces the highest average *p_BOLD_*, compared to all other pre-processing pipelines. We also show that adding the global signal as a regressor increases TSNR but decreases *p_BOLD_*. By looking at the BOLD properties of the global signal (GS) and the amount of variance from the GS explained by cardiac and respiratory regressors, we conclude that the observed reduction in *p_BOLD_* is likely due to the removal of neurally-driven BOLD fluctuations.

Finally, to assess if higher *p_BOLD_* results in better data from an application perspective, we attempted prediction of fluid IQ scores using connectome predictive modeling (CPM; (Shen et al., 2017). We observe that prediction accuracy and precision showcase the same across-pipeline profile as *p*_*BOLD*,_ and not that of TSNR. In other words, *p_BOLD_* forecasts the ability to predict fluid IQ better than TSNR.

In summary, this work introduces, evaluates and demonstrates a novel QA metric for ME data. Our results suggest that this new metric outperforms TSNR in three application scenarios. Additionally, using p_BOLD_ we show that GS regression (GSR) lowers BOLD contents, and results in less precise prediction of fluid IQ. Regarding this topic, our findings support literature suggesting that the GS is not necessarily dominated by motion and respiratory artifacts, but that it also contains a significant amount of information regarding neuronal-related processes (Liu et al., 2017).

## Theory

### Signal Model

Assuming a mono-exponential decay, the gradient recalled echo (GRE) signal at a voxel *x*, for time *t*, and echo time *TE*_*i*_ is given by:

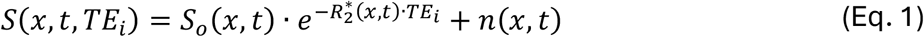

where *S*_*o*_(*x, t*) denotes fluctuations in net magnetization (i.e., non-BOLD), 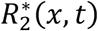 denotes changes in transverse relaxation rate (i.e., 1/T_2_^*^ including BOLD) and *n*(*x, t*) is the noise term.

When working with ME data, it is often preferable to write the net magnetization (*S*_*o*_(*x, t*)) and transverse relaxation 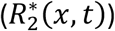 contributions in terms of their temporal means 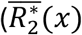 and 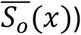 and their respective change with respect to that mean 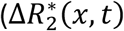 and Δ*S*_*o*_(*x, t*))

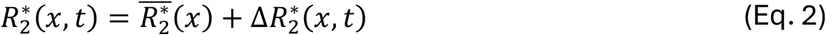

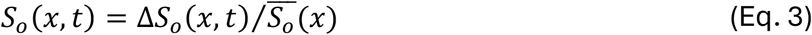

When the noise term is negligible, the measured signal in units of signal percent change (*S*_*spc*_) can be approximated linearly as follows:

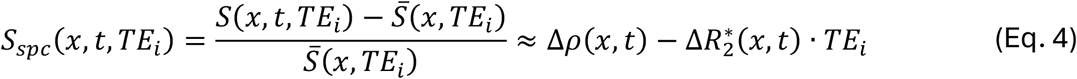

where Δ*ρ*(*x, t*) is defined as: 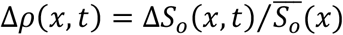. For simplicity, we can then rewrite Eq. (4) as follows:

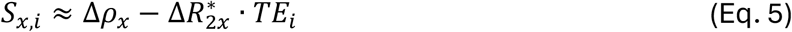

Equation 5 then represents the measured signal over time at a given location *x* for the echo time *TE*_*i*_ in units of signal percent change.

### Across-echoes Functional Connectivity using Pearson’s correlation

In fMRI, the most common way to estimate synchronicity (or functional connectivity (FC)) between recordings at two different voxels or regions of interest (ROIs) is using Pearson’s correlation (*r*). If we consider signals from locations *x* and *y* that were measured at echo times *TE*_*i*_ and *TE*_*j*_, respectively, their Pearson’s correlation coefficient, or *FC*_*r*_ in the following, is given by:

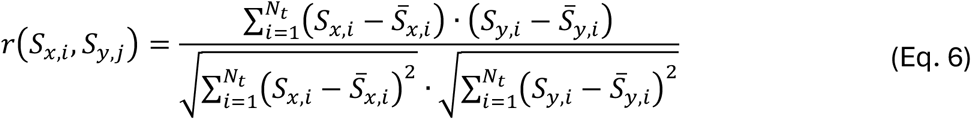

where *N*_*t*_ is the number of acquisitions. Hereinafter, the summatory term is denoted as Σ. When the signals are given in units of signal percent change, their means equal zero 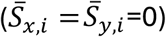, and Eq. 6 can be simplified as:

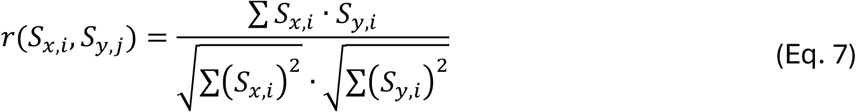

If we now substitute these signals by their formulation in Eq. (5), the across-echo *FC*_*r*_ becomes:

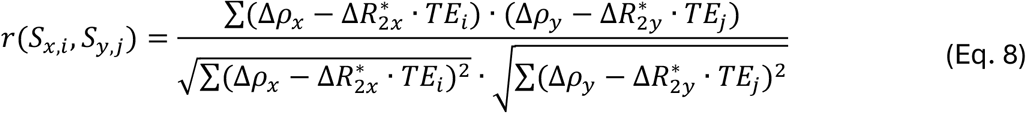

Using equation 8 we can now explore how FC across-echoes behaves in two extreme signal regimes:

1. <u>Data dominated by S_o_ fluctuations 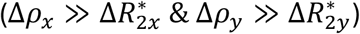:</u> In this case, the across-echo *FC*_*r*_ can be approximated as:

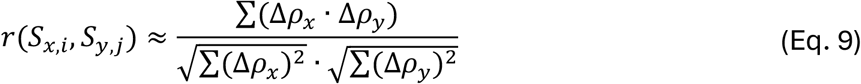

This shows the FC estimated using Pearson’s correlation is independent of the TEs when the data is dominated by S_o_ fluctuations.
2. <u>Data dominated by BOLD fluctuations 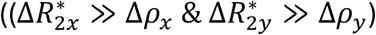:</u> In this case, the across-echo *FC*_*r*_ can be approximated as:

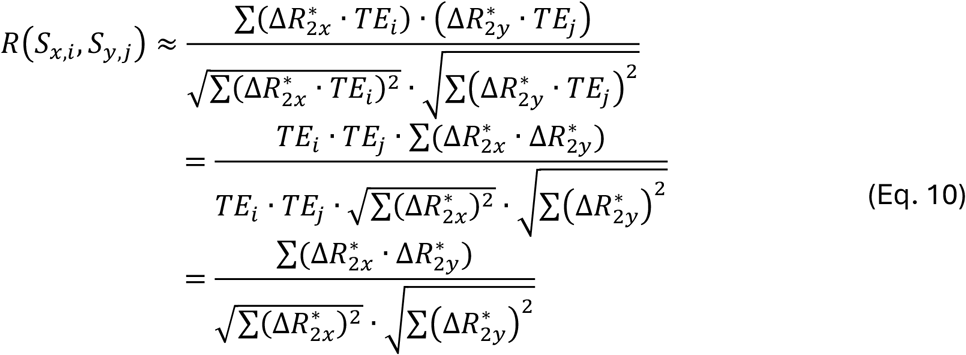

This shows that when BOLD fluctuations dominate, *FC*_*r*_ is also echo-time independent. In summary, FC estimated with Pearson’s correlation is TE-independent irrespective of whether BOLD or S_o_ fluctuations dominate the recordings.

### Across-echoes Functional Connectivity using Covariance

An alternative way to quantify inter-regional FC is using the covariance (*C*) between the signals, which is defined as:

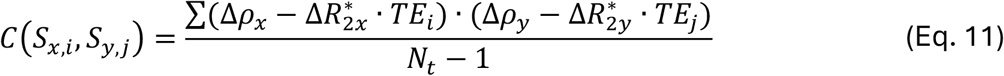

For the two extreme regimes described above, we now observe differing behaviors:

1. <u>Data dominated by S_o_ fluctuations 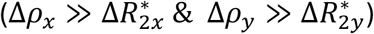</u>: In this case, Eq. 11 can be simplified as follows:

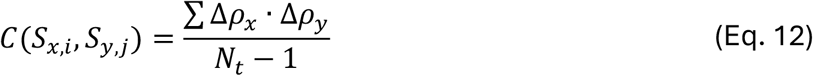

This shows that covariance-based FC (*FC*_*C*_) is TE-independent when the data is dominated by S_o_ fluctuations.
2. <u>Data dominated by BOLD fluctuations 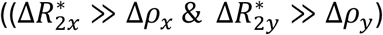</u>: In this case, Eq. 11 becomes:

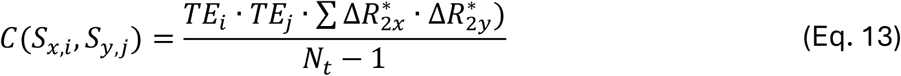

This shows that covariance-based FC is TE-dependent if the data is dominated by BOLD fluctuations.

In summary, covariance-based FC shows two clearly distinct behaviors depending on whether the data are dominated by BOLD or S_o_ fluctuations. We can now extrapolate this observation made for two locations (*x* and *y*) to full brain FC matrices based on *N* regions of interest (ROIs). For that, we can visualize scatter plots of FC computed using one pair of echo times (e.g., (*TE*_*i*_, *TE*_*j*_)) against FC computed using a second pair of echo times (e.g., (*TE*_*k*_, *TE*_*l*_)). In such scatter plots, edges (or connections) are represented by points whose coordinates are dictated by the strength of FC estimated using the corresponding TEs. Figure 1a illustrates a simulation of how such a scatter plot would appear when data are S_o_ dominated: As covariance-based FC (*FC*_*C*_) is TE-independent, edges are expected to lay relatively close to the identity line (dashed red line). Figure 1b shows a representative example based on cardiac-gated data, which is expected to be dominated by S_o_ fluctuations. We observe that edges lay near the identity line for this scan.

**Figure 1.**
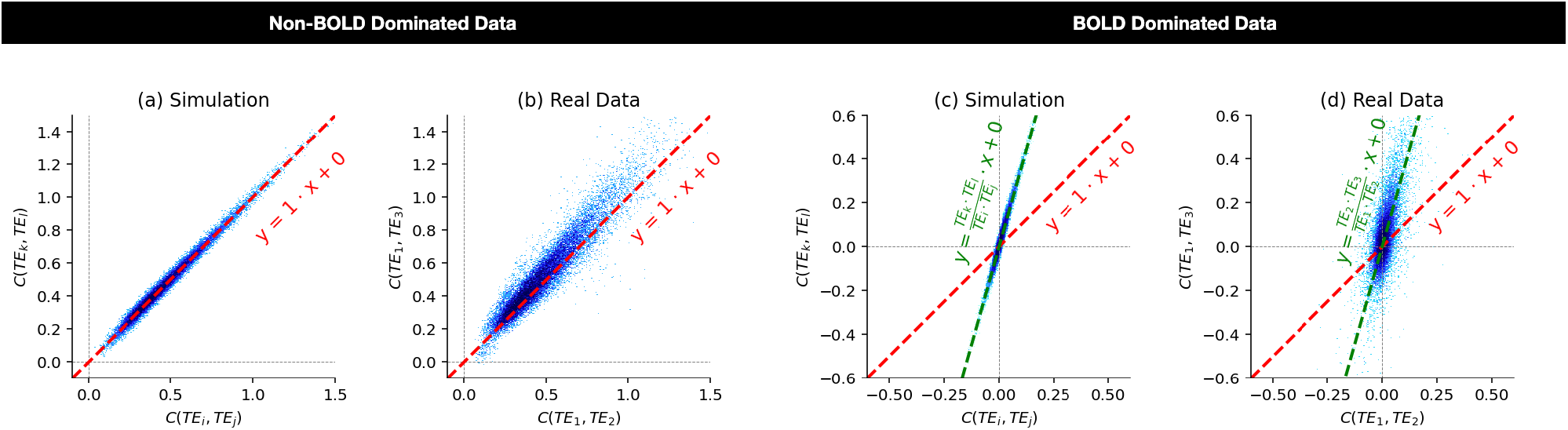
Schematic (a,c) and real-data examples (b,d) of expected behaviors for across-echoes covariance-based FC matrices in two extreme signal regimes: non-BOLD dominated and BOLD dominated data. (a) Schematic of expected behavior for non-BOLD dominated data. In this case, edges sit in or near the identity line (red dashed line). (b) Same as (a) but using a representative cardiac-gated dataset with minimal denoising. (c) Schematic of expected behavior for BOLD dominated data. Here, edges sit primarily over a line of slope dictated by contributing echoes (green dashed line) as opposed to the identity line (red dashed line). (d) Same as (d) but using a representative low motion, constant-gated scan following tedana denoising. This figure was created using notebook N04a_Figure01_Simulation.ipynb available in the github repo that accompany this publication (see code availability section below).

Conversely, Figure 1c shows a simulation of expected behavior when data is BOLD dominated where *FC*_*C*_ is expected to be TE-dependent. In this case, edges no longer lay over the identity line (red dashed line), but over a line with intercept equal to zero and slope given by the ratio of the contributing TEs (green dashed line). Figure 1d shows an example with low-motion, fully pre-processed constant-gated data. We observe that edges lay closer to the line with the TE-dependent slope (green dashed line) than to the identity line (red dashed line).

### p_BOLD_: a new QC Metric for Multi-Echo Data

Based on the theory just discussed, we propose a new QC metric for ME data called *p_BOLD_* (short for probability of data being BOLD dominated). This metric quantifies how well the data sit near the line with TE-dependent slope as opposed to near the identity line. One can quantify these behaviors in different ways. Perhaps the most direct approach would be by fitting a linear model to the cloud of points representing the edges, and then comparing the slope and intercept of this linear model to those of the lines associated with BOLD and S_o_ dominated regimes. While conceptually simple, this procedure is problematic because fMRI data are often heteroscedastic, and a linear fit produces biased estimates of slope and intercept. Supplementary Figure 1a-b exemplifies this issue with one representative scan from the evaluation dataset. Supplementary Figure 1a shows in red the identity line (S_o_ behavior), in green the line associated with BOLD dominated data, and in blue the linear fit to the data. Although edges fall primarily over the green line, the linear fit (blue line) does not. Supplementary Fig 1b shows the residual plot for the fit. Here, one can clearly observe that the spread of residuals is not uniform across the range of fitted values. In fact, the heteroscedasticity of this sample scan can be further confirmed statistically using the Breusch-Pagan Lagrande Test (1979), BPL stat = 1790 (*α* < 0.05).

To avoid this issue, we propose to estimate *p_BOLD_* using the average preference of edges towards each of the two lines of interest (BOLD and S_o_ line). Given a ME dataset with *N*_*e*_ echoes 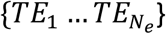 and a brain parcellation with *N*_*r*_ ROIs {*x, y, z*, … }, *p_BOLD_* is computed with the following steps (Figure 2):

1. Compute the *FC*_*c*_ matrix for a pair of echo times (*TE*_*i*_, *TE*_*j*_): *FC*_*C*_[(*x, y*), *TE*_*i*_, *TE*_*j*_] (Fig. 2a).
2. Compute the *FC*_*c*_ matrix for a second pair of echo times: *FC*_*C*_[(*x, y*), *TE*_*k*_, *TE*_*l*_] (Fig. 2b).
3. Take the top triangle of these two matrices to map all edges into a 2D plane. Here, each edge (*x, y*) is represented by a black point with the abscissa given by *FC*_*c*_[(*x, y*), *TE*_*i*_, *TE*_*j*_] and the ordinate given by *FC*_*C*_[(*x, y*), *TE*_*k*_, *TE*_*l*_] (Fig. 2c).
4. For each edge (*x, y*) estimate 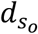: the distance between the 2D point representing the edge and the line representing S_o_ dominated behavior (i.e., the identity line). To estimate 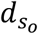, one can first project the edge (black point in Fig. 2d) to the S_o_ line (red dashed in Fig. 2d) and then compute the length of the segment that joins these two points (red continuous line in Fig. 2d). Fig. 2e shows edges colored by 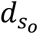 for a representative scan.
5. Similarly, for each edge (*x, y*), estimate *d*_*BOLD*_: the distance between the 2D point representing the edge and the line representing BOLD dominated behavior (i.e., green dashed line with zero intercept and slope equal to *TE*_*k*_ · *TE*_*l*_/*TE*_*i*_ · *TE*_*j*_). Fig. 2f shows the process for a representative edge, and Fig. 2g shows all edges colored by *d*_*BOLD*_.
6. Given 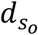 and *d*_*BOLD*_, and a tolerance threshold (*δ*), the preference of each connection towards the BOLD line can be labelled as follows:

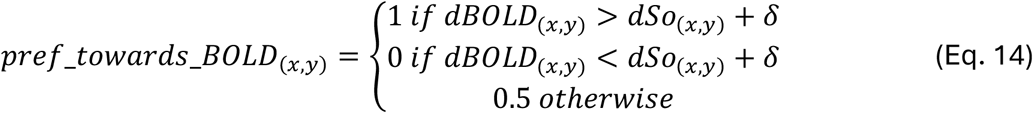

In this work, the tolerance was set to *δ* = 10^−3^. Figure 2h shows all edges colored according to their preference value.
7. Because edges laying close to the origin have more similar distances to both target lines than those further away from the origin, we weight edge-wise preference values towards the BOLD line using the distance of the edges to the origin:

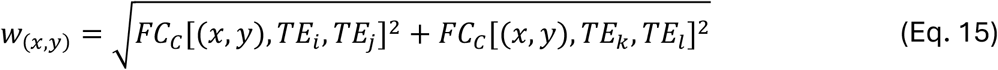

Fig. 2i shows edges colored by weights, and Fig. 2j shows connections colored by their weighted preference towards the BOLD line.
8. The *p_BOLD_* for a quadruple of echo pairs ((*TE*_*i*_, *TE*_*j*_),( *TE*_*k*_, *TE*_*l*_)) is given by the weighted average of edge-wise preference towards the BOLD line:

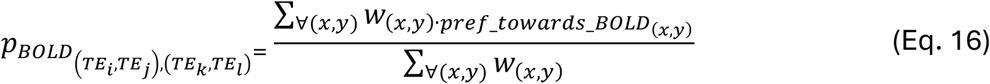
9. To obtain a final, single, *p_BOLD_* value per scan, we repeat steps 1 – 8 using all possible combinations of echoes (e.g., *[(TE*_*1*_,*TE*_*1*_*),(TE*_*2*_,*TE*_*2*_*)], [(TE*_*1*_,*TE*_*2*_*),(TE*_*2*_,*TE*_*2*_)], etc.); and then compute the average. Because the separation between the BOLD and S_o_ lines depends on the contributing TEs—and so does our ability to discern edge preference; during this final average step, we weight individual p_BOLD_ estimates by the chord distance between the two lines for each respective TE quadruple as follows:

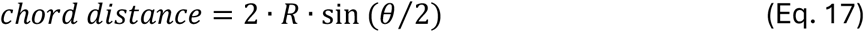

where R denotes the radius of an intersecting circumference centered at the origin (R=0.5 in this work) and *θ* denotes the angle between both lines. Figures 2k-l depict two examples for two different TE quadruples.

**Figure 2.**
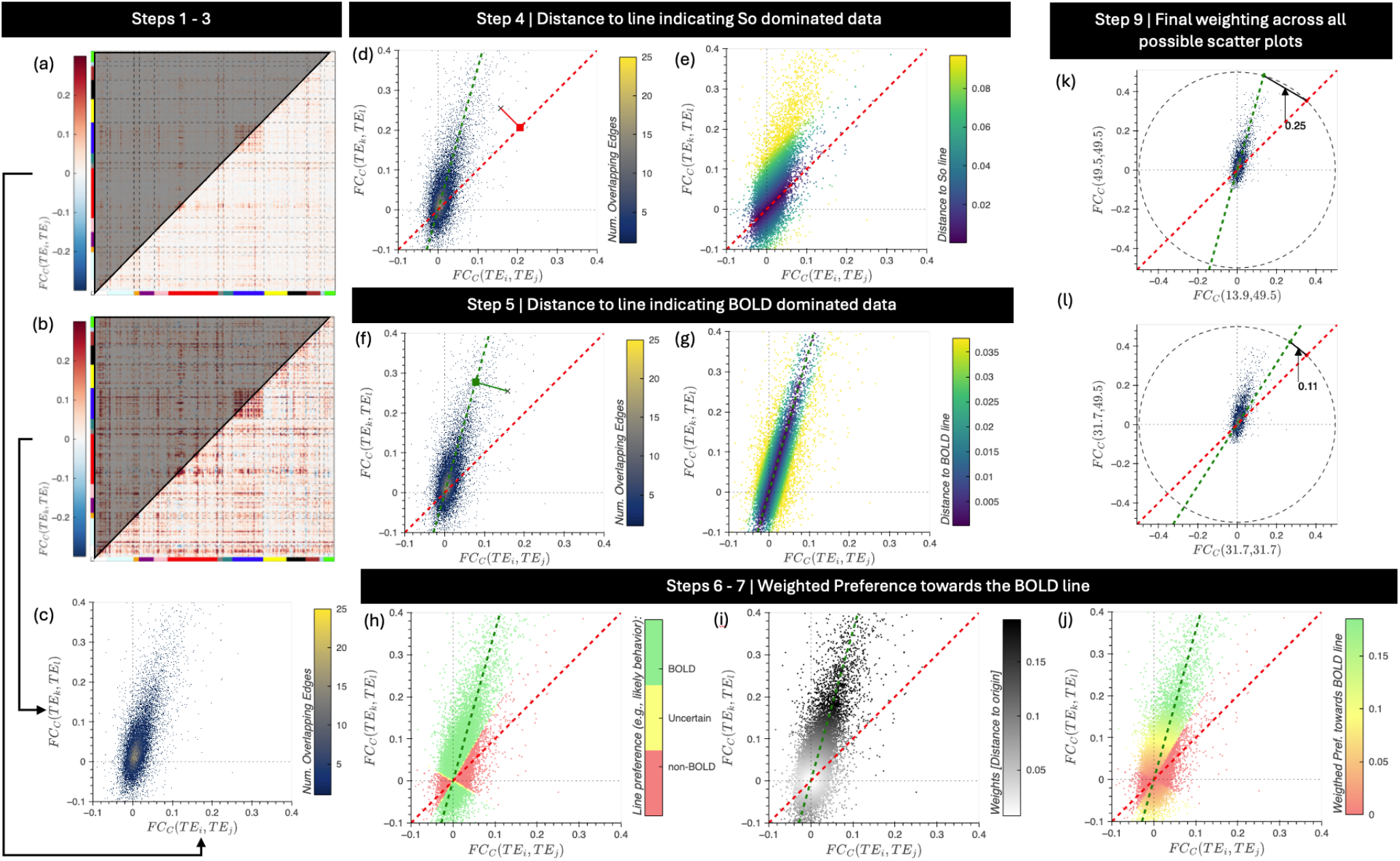
Steps involved on the estimation of the p_BOLD_ quality assurance metric. (a) Covariance matrix computed using a representative pair of echo times (TE_i_,TE_j_). (b) Covariance matrix calculated using a second pair of echo times (TE_k_,TE_l_). (c) Scatter plot of edge covariances computed with two different pairs of echo times. In this plot each edge is represented by a dot. To highlight the high density of connections near the origin, we color the plot as a function of density of points. (d) Example of how to compute the distance between a given edge (black X) and a line representing data dominated by So fluctuations (red dashed line). (e) Scatter plot with edges (e.g., points) colored according to their distance to the So line. (f) Example of how to compute the distance of an edge to the BOLD line (green dashed line). (g) Scatter plot with edges (e.g., points) colored according to their distance to the BOLD line. (h) Scatter plot with edges colored according to their preference towards the BOLD line (green = 1; yellow = 0.5, red = 0) as defined in Eq. 14. (i) Scatter plot with edges colored according to their distance to the origin, which will be used during averaging. (j) Scatter plot of edges colored according to the weighted preference towards the BOLD line. We can observe how the contribution of preference values associated with edges near the origin is diminished by this weighting scheme. (k-l). Example of BOLD and So lines for two different echo time quadruples. In each case, we show the chord distance between the lines computed for a radius of 0.5.

## Methods

### Datasets

This work uses two different datasets to validate and apply the proposed *p_BOLD_* metric. First, we use a small (n=7) dataset—labelled the “discovery” dataset—to empirically demonstrate the theoretical observations described in the theory section. Second, we use a large (n=439) dataset—labelled the “evaluation dataset”—to further support our theory on a larger additional sample, and also to exemplify how *p_BOLD_* can be used effectively when working with large-N datasets.

#### Discovery Dataset

Two resting-state scans were acquired in 7 different participants using a General Electric (GE) 3 Tesla 750 MRI scanner (Waukesha, WI). The scanner’s body coil was used for RF transmission, and a 32-channel receive-only head coil (GE, Waukesha, WI) was used for reception.

Resting-state scans were acquired with a multi-echo EPI sequence (flip angle = 60°, TEs = 13.9/31.7/49.5 ms, 33 oblique slices, slice thickness = 3 mm, in-plane resolution=3×3 mm^2^, acceleration factor=2, number of acquisitions = 195, bottom/up sequential acquisitions). For each participant, we acquired one non-gated functional scan and one cardiac-gated functional scan. Non-gated scans were acquired using a constant repetition time (TR) of 2.5 s. Cardiac-gated acquisitions were time-locked to the first peak of the cardiac cycle, recorded on a GE optical pulse oximeter attached to one of the subject’s fingers, following a nominal TR of 2.5 s. This resulted in a non-constant TR of mean ± SD = 3.3 ± 0.7 s across the dataset (one scan excluded from this computation because trigger file was not saved).

In addition, *T1*-weighted Magnetization-Prepared Rapid Gradient-Echo (MPRAGE) and Proton Density (PD) sequences were acquired for presentation and alignment purposes (number of slices per slab= 176; slice thickness= 1 mm; matrix size= 256 × 256).

In addition to the resting-state scans and the anatomical scans, task-based scans were also acquired for other purposes. Those are described in (Gonzalez-Castillo et al., 2016). This data was collected after obtaining informed consent in compliance with the NIH Combined Neuroscience Institutional Review Board (approved protocol 93-M-0170) in Bethesda, MD.

#### Evaluation Dataset

The evaluation dataset consists of 602 resting-state scans from the publicly available Neurocognitive Aging Dataset collected by Spreng et al.(2022). Full details about this dataset can be found on the Scientific Data publication that accompanied the data release. Here, we provide acquisition information regarding the data acquired on site 1 (Cornell Magnetic Resonance Imaging Facility); as this is the data used in this work.

Three hundred and one participants completed two resting-state scans (duration: 10 mins and 6 secs) on a GE 3T scanner. Scans were acquired using a multi-echo sequence with the following parameters (TR = 3 s, TE=13.7/30/47 ms, flip angle = 87º, matrix size = 72×72, 46 axial slices, 3 mm isotropic voxels, 204 volumes, 2.5x acceleration with sensitivity encoding). For 233 participants, pulse and respiration were monitored continuously during scanning using an integrated pulse oximeter and respiratory belt.

Additionally, T1 anatomical scans were acquired using a T1-weighted volumetric magnetization prepared rapid gradient echo sequence (TR = 2530 ms; TE = 3.4 ms; 7° flip angle; 1 mm isotropic voxels, 176 slices, matrix size=256×256) with 2x acceleration with sensitivity encoding.

Of the original 602 resting state scans in the Spreng dataset, only 476 were acquired on Site 1. Of these, only 439 scans entered the analyses. Table 1 summarizes exclusion criteria and the number of scans affected.

**TABLE 1.** Scan exclusion criteria and scan count entering the analyses.

| *Elimination Criteria* | *Scans Removed* | *Scans Remaining* |
| --- | --- | --- |
| *Failed functional to anatomical alignment* | 19 | 457 |
| *Voxel size differs* | 16 | 441 |
| *Number of acquisitions differs* | 2 | 439 |

Finally, this data release also includes a series of behavioral and phenotypic metrics for a large portion of scanned subjects. Here, we will use the NIH fluid composite IQ score to exemplify how well *p_BOLD_* forecasts the ability to predict phenotypic scores based on FC.

### Data Pre-processing

High resolution T1 anatomical scans were first submitted to Freesurfer (Fischl, 2012) *recon-all* to create skull-stripped, intensity corrected versions of the anatomical scan; and also subject specific binary masks of the grey matter (GM) ribbon, white matter (WM) and lateral ventricles. All these outcomes were used in subsequent analyses as described below.

Functional data were pre-processed using three different pipelines that differed in terms of the nuisance signals used during the final regression step. Prior to regression, all pipelines shared the following pre-processing steps: discard initial three volumes, despiking, time shift correction, head motion estimation, and non-linear corregistration to MNI space (using AFNI provided template: *MNI152_2009_template_SSW*). All these pre-processing steps were performed with AFNI’s *afni_proc.py* (Reynolds et al., 2024). The next step in all pipelines was nuisance regression. Nuisance regression was separately applied to the data at each echo time. These are the signals regressed in each pipeline:

- <u>Basic Pipeline [Basic]:</u> This pipeline includes as nuisance regressors the six head motion estimates, their first temporal derivatives, Legendre polynomials up to 5^th^ order to remove slow signal drifts, *aCompCorr* physiological regressors (signal from WM plus 3 principal components from the ventricular mask; (Behzadi et al., 2007)), and sines and cosines needed to model frequency components outside the 0.01 – 0.2 Hz range.
- <u>Global Signal Regression Pipeline [GSR]:</u> In addition to the regressors included in the Basic pipeline, this pipeline also includes the global signal (GS). The GS was computed separately for each TE using a full brain mask generated by AFNI and then restricted to voxels with signal in all echoes, according to tedana (DuPre et al., 2021). We use this mask because it ensures we only consider voxels free of significant signal dropout in all TEs.
- <u>Tedana Denoising Pipeline [Tedana]:</u> In this case, in addition to the Basic regressors we also remove signals from components marked as “*unlikely BOLD*” by tedana (DuPre et al., 2021). This was accomplished using tedana (regression of “unlikely BOLD components) followed by AFNI *3dTproject (*regression of Basic nuisance signals). This procedure mimics AFNI *afni_proc*’s denoising procedure for multi-echo data, except that here we work with the individual echoes as opposed with the optimally combined timeseries.

All three pipelines were also run on data that underwent magnitude-only NORDIC (m-NORDIC; (Moeller et al., 2021)) denoising prior to entering any pre-processing; leading to a total of six different sets of fully pre-processed data to be compared.

### Estimation of Temporal Signal-to-Noise Ratio

One of the most prevalent QA metrics in fMRI is the Temporal Signal-to-Noise Ratio (TSNR), which is defined voxel-wise as the mean across time divided by the standard deviation across time (Parrish et al., 2000). TSNR maps were estimated following the regression step in each of the six pre-processing pipelines just described. Because all pipelines include regressors to remove slow signal drifts, the mean signal was reintroduced onto the residuals prior to TSNR computation. As such, TSNR is always computed on detrended data. We then used AFNI program *compute_ROI_stats.tcsh* to compute a single representative TSNR value per scan: the median TSNR across all voxels inside the brain. We only report TSNR for the second TE.

### Estimation of Functional Connectivity Matrices

Functional connectivity matrices, whether based on Pearson’s correlation (*FC*_*R*_) or Covariance (*FC*_*C*_), were estimated using the “*Powers 264 ROI*” brain parcellation (Power et al., 2011). For each dataset, we only included ROIs that were at least 95% inside the imaging field of view for all scans in the dataset. This restricted the analyses in the discovery dataset to 226 ROIs, and 203 ROIs for the evaluation dataset. Discarded ROIs fall primarily within inferior brain regions with large signal dropout or ROIs sitting near the lower edge of the imaging field of view.

Using the final parcellations, we estimated representative ROI timeseries—namely the spatial mean across all voxels in each ROI—for each TE separately. We then computed functional connectivity matrices, *FC*_*R*_ and *FC*_*C*_, using all 15 possible echo times quadruples (e.g., *[(TE*_*1*_,*TE*_*1*_*),(TE*_*1*_,*TE*_*2*_*)], [(TE*_*1*_,*TE*_*1*_*),(TE*_*1*_,*TE*_*3*_*)]… [(TE*_*2*_,*TE*_*3*_*),(TE*_*3*_,*TE*_*3*_)]) given the three available echo times (*TE*_*1*_, *TE*_*2*_, *TE*_*3*_).

### Estimation of *p_BOLD_*

For each pre-processing pipeline, we estimated p_BOLD_ (steps 3-8 above) for each of the 15 unique echo time quadruples available. Then, we obtained a single *p_BOLD_* value per scan by computing a weighted average of these *p_BOLD_* values (step 9 above).

### Thermal Noise Estimation

NORDIC denoising (Moeller et al., 2021) is intended to remove thermal noise from fMRI scans prior to any subsequent pre-processing. To evaluate the effectiveness of m-NORDIC in our dataset, we estimated thermal noise separately on each echo before and after m-NORDIC.

Optimal thermal noise estimation requires the acquisition of noise scans with the same readout gradients, but no excitation pulse. When such scans are not collected, as it is the case here, an approximate estimate of thermal noise can be computed as the spatial standard deviation of the unprocessed data in image space on object- and ghosting-free voxels (Chang et al., 2005). In this work, we define object- and ghosting-free voxels as those with mean signal levels of 2 or less in all TEs. To ensure estimates are not biased by voxels with null recordings, we exclude voxels with zero counts above 20.

m-NORDIC reduced thermal noise by approx. 31.6% +/- 4.3% in the discovery dataset and 28.8% +/- 1.8% in evaluation dataset. These reductions were statistically significant according to paired T-tests (p<1e^-4^) conducted separately for each echo time (See Suppl. Figure 2 for details).

### Estimation of Kappa and Rho for the Global Signal

Echo time-dependent denoising (Evans et al., 2015; Kundu et al., 2014), as implemented in tedana, calculates two metrics—*kappa* and *rho*— as respective measures of contributions of T_2_^*^ and S_o_ fluctuations in each ICA component. To calculate the T_2_^*^ and S_o_ contributions to the GS, use the tedana function ‘*generate_metrics*’ to estimate the *kappa* and the *rho* of the GS. To do so, we provide the GS computed on the optimally combined timeseries as input to this function.

### Variance Explained by Physiological Noise in the Global Signal

To evaluate the contribution of cardiac and respiratory effects to the GS, we calculated the amount of variance in the GS that could be explained by common physiological regressors.

This portion of the analyses only included 370 scans for which complete accompanying cardiac and respiratory traces were available. For these scans, we first estimated cardiac and physiological regressors associated with the RETROICOR (Glover et al., 2000) and RVT (Birn et al., 2006) models using AFNI program ‘*physio_calc.py*’. This program generates 13 physiological regressors: eight correspond to RETROICOR-based regressors and five to RVT-based regressors. Because the primary goal ‘*physio_calc.py*’ is to create regressors for denoising volumetric fMRI data, the program generates one time-shifted version of each of these 13 regressors per slice. In our data, this translates to a total of 598 regressors (46 slices x 13 regressors); which is well above the number of available repetitions. Because the objective here is to estimate how much variance can be explained by physiological traces in a single recording—namely the GS—for each of the 13 regressors, we select the time-shifted regressors showing the largest correlation with the GS.

Finally, we use ordinary least squares (OLS) to estimate the adjusted R^2^ (our measure of explained variance) of a model containing these 13 physiological regressors as explanatory variables and the GS as the dependent variable. We perform this operation using recordings from the second echo time, as this is the one with the highest BOLD contrast. The OLS model and adjusted R^2^ (adjR^2^) estimation was performed using the *statsmodel* python library. Statistical significance of adjR^2^ estimates was evaluated against 10,000 random permutations in which GS and physiological regressors came from randomly selected scans.

### Prediction of Fluid Cognition Composite Score (fluid IQ)

One common application of fMRI, specially resting-state data, is to find associations between functional connectomes and phenotypes. A practice often known as brain-wide associations. To evaluate the informative value of p_BOLD_ in this context, we attempt prediction of the NIH Fluid Cognition composite cognitive score (fluid IQ; (Akshoomoff et al., 2013)) of the subjects in the evaluation dataset. The ability to predict it using FC is well stablished in the literature (Dubois et al., 2018; Finn et al., 2014; Wilcox and Barbey, 2023).

For this portion of the work, we used the connectome-based predictive modeling algorithm (CPM; (Shen et al., 2017)). To mitigate the effect of head motion confounds, we residualized the target variable (i.e., fluid IQ) with respect to mean head motion before training the models. We ran CPM separately for each scanning session. We used data from 200 subjects from session 1 and 201 subjects from session 2 for whom fluid IQ scores are available.

We ran the CPM algorithm one hundred times per session to avoid any bias due to random assignment of scans to testing and training folds. On each such iteration, data was randomly divided into training and test sets using a 10-fold cross validation procedure. For feature selection, we used Spearman’s correlation and a p-value threshold of 0.01. We report accuracy for the composite model that considers edges that positively and negatively correlate with the prediction target. Prediction is quantified as the Pearson’s correlation between observed and predicted values.

## Results

### Discovery Dataset

The discovery dataset provides empirical data that mimics the two extreme scenarios described in the theory section. First, cardiac-gated data pre-processed using the Basic pipeline without NORDIC denoising delivers data dominated by S_o_ effects because of the irregular sampling time associated with cardiac-gated acquisitions. Second, constant-gated, low motion, tedana pre-processed data with m-NORDIC denoising offers high quality data dominated by BOLD fluctuations. Figure 3a shows how *p_BOLD_* was able to effectively capture this distinction in this dataset: low *p_BOLD_* for S_o_-dominated and high *p_BOLD_* for BOLD-dominated data. Figure 3b shows *FC*_*R*_ matrices for the two fully pre-processed, constant-gated scans with the highest (*p_BOLD_*=0.92) and lowest (*p_BOLD_*=0.85) *p_BOLD_* values. These matrices were computed using the second TE. In both cases, we see a stereotypical FC structure with clear delineation of networks around the diagonal. Figure 3d shows scatter plots computed using the TE quadruple *[(TE*_*1*_,*TE*_*1*_*),(TE*_*3*_,*TE*_*3*_)] for the same two datasets. Here we can observe how connections concentrate around the line associated with BOLD-like behavior (dashed green line) and away from the line associated with S_o_-like behavior (dashed red line). Figures 3c and 3e show two representative scans from the cardiac-gated sample with minimal pre-processing (i.e., Basic pipeline without m-NORDIC). For the scan with the lowest *p_BOLD_* (*p_BOLD_*=0.04), we observe an FC matrix with little structure and widespread high correlation values. This is due to the ubiquitous presence of S_o_ fluctuations caused by the irregular TR contaminating all ROIs. The second cardiac-gated scan presented on the left of Figure 3c corresponds to the outlier cardiac-gated scan with the highest *p_BOLD_* (*p_BOLD_*=0.77). This high *p_BOLD_* value suggests that this particular scan is not dominated by S_o_ fluctuations, despite being acquired with an irregular TR. The *FC*_*R*_matrix associated with this scan (right of Fig 3c) shows clear network structure and the scatter plot below shows edges that fall primarily over the BOLD-like line. This suggests that S_o_ fluctuations are not dominant in this scan; which could only result if TR variability across acquisitions was low. Figure 3f confirms that is the case. For the scan with *p_BOLD_*=0.04 we see a wide spread of TR offsets (light orange distribution), while for the scan with *p_BOLD_*=0.77, most TR delays are narrowly centered around 900 ms.

**Figure 3.**
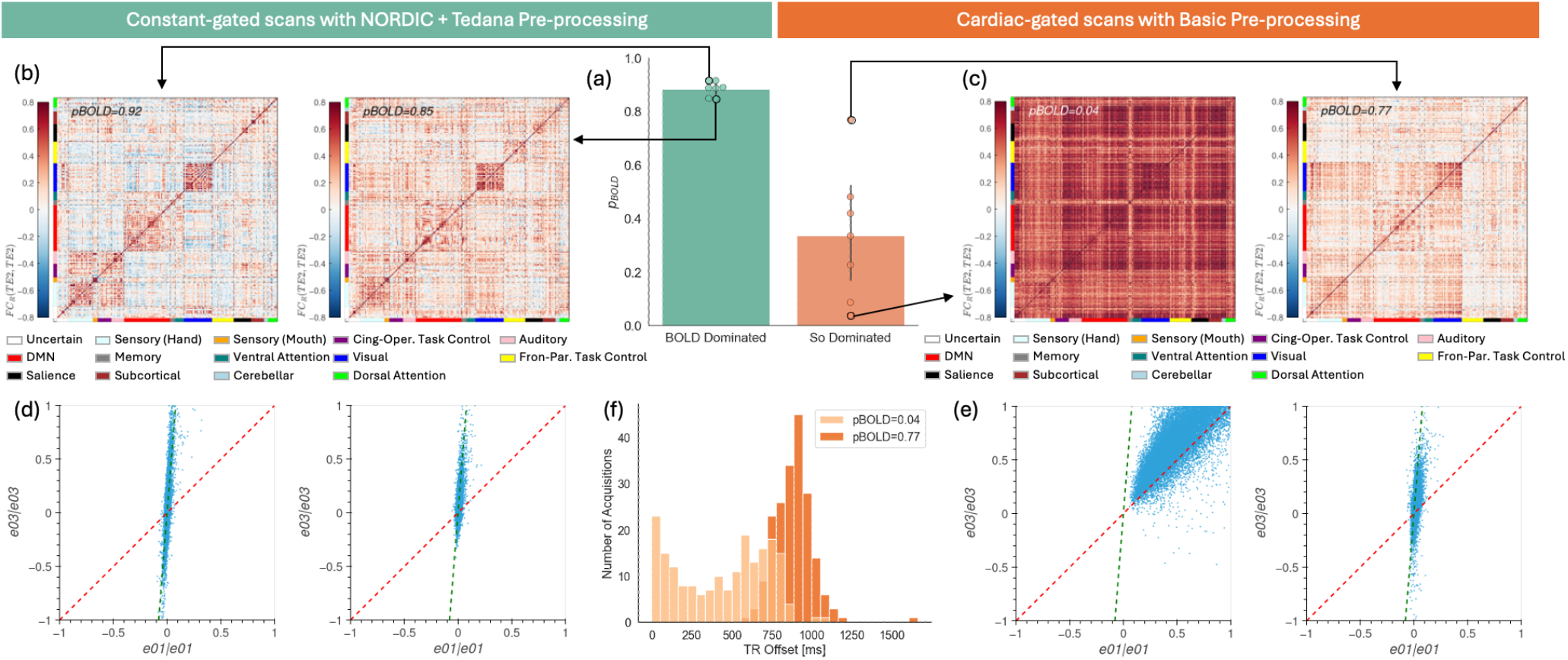
Exploration of p_BOLD_ behavior in the discovery dataset (a) Bar plots showing group average (as bars) and individual p_BOLD_ values (as dots) for data expected to be dominated by So fluctuations (orange) and BOLD fluctuations (light green). (b) Person’s based FC matrices for the constant -gated scans with highest and lowest p_BOLD_. (c) Pearson’s based FC matrices for the cardiac-gated scans with highest and lowest p_BOLD_. (d) Scatter plots of covariance FC for the [(TE_1_,TE_1_),(TE_3_,TE_3_)] echo times quadruple for the same two scans shown in (b). Individual edges are represented as points. The red dash line is the identity line representing So-dominated behavior. The dashed green line represents BOLD-dominated behavior. (e) Same as (d) but for the cardiac-gated scans shown in (c). (f) Distribution of acquisition delays relative to the nominal TR of 2.5s for the two cardiac-gated scans shown in (c).

### Evaluation Dataset

Once we confirmed that *p_BOLD_* follow theoretical expectations on the discovery dataset, we moved to the evaluation dataset to understand how *p_BOLD_* can provide meaningful information beyond that available using traditional QA metrics such as TSNR.

#### Pre-processing Pipeline Evaluation with p_BOLD_ and TSNR

TSNR significantly increased for GSR pipeline relative to the Basic pre-processing pipeline and for the tedana pipeline relative to the GS and Basic pipelines. Adding m-NORDIC also increased TSNR for each of the three pipelines. Incremental gains associated with tedana are greater when m-NORDIC is applied. (Figure 4a, Table 2).

**TABLE 2.** Mean, Standard Deviation and Percent Change for TSNR and pBOLD when m-NORDIC is on and off in each pipeline. Data corresponds to all scans in the evaluation dataset.

| Pipeline | Basic |  |  | GS Regression |  |  | Tedana |  |  |
| --- | --- | --- | --- | --- | --- | --- | --- | --- | --- |
| m-NORDIC | Off | On | % Increase | Off | On | % Increase | Off | On | % Increase |
| TSNR | 122±14 | 170±31 | 28% | 126±14 | 179±32 | 30% | 143±13 | 247±46 | 42% |
| $p_{BOLD}$ | 0.82±0.08 | 0.83±0.09 | 1.2% | 0.78±0.08 | 0.79±0.09 | 1.3% | 0.86±0.04 | 0.87±0.04 | 1.1% |

**Figure 4.**
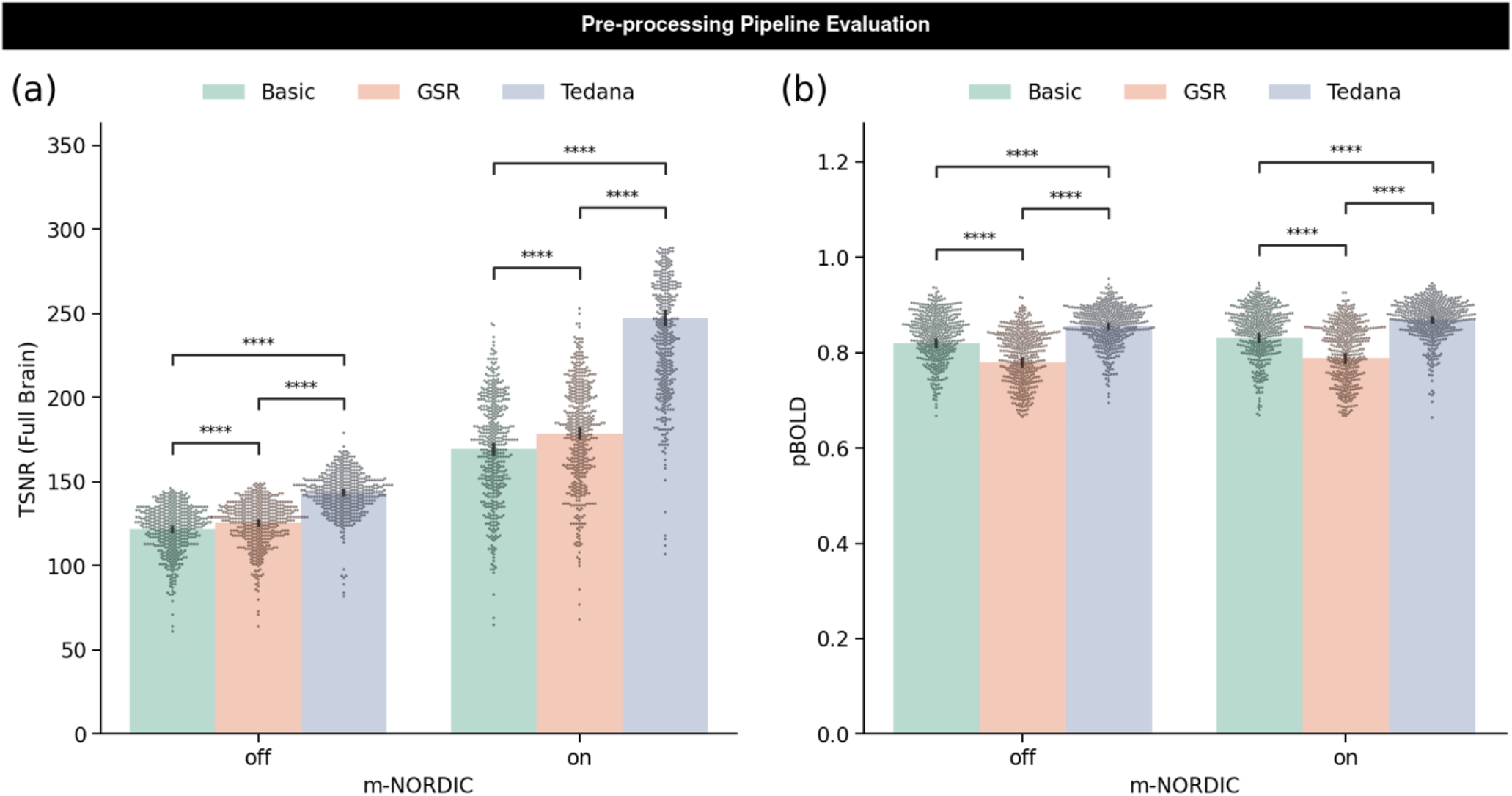
Evaluation of pre-processing pipelines using TSNR and p_BOLD_. These results correspond to all scans in the evaluation dataset (a) TSNR for all six pre-processing pipelines. Bars represent the mean and black lines the 95% confidence interval. Individual dots represent scans. Statistical significance estimated using the Mann Whitney test (**** p<10^-5^). (b) p_BOLD_ for all six pre-processing pipelines.

Similarly to TNSR, tedana results in significant increases in *p_BOLD_* relative to the Basic pipeline, but, unlike TSNR, GSR does not (Figure 4b, Table 2). Indeed, GSR results in lower *p_BOLD_* than Basic pre-processing. This is true for both m-NORDIC scenarios.

#### Identification of problematic scans with p_BOLD_ and TSNR

Next, figure 5a shows a scatter plot with each scan represented by a 2D point with the x-coordinate given by its *p_BOLD_* and the y-coordinate by its TSNR. This scatter plot was constructed using data from the Basic pipeline without m-NORDIC. The goal here is to exemplify how, given a pre-processing pipeline, *p_BOLD_* and TSNR can provide complementary information regarding individual scan quality.

**Figure 5.**
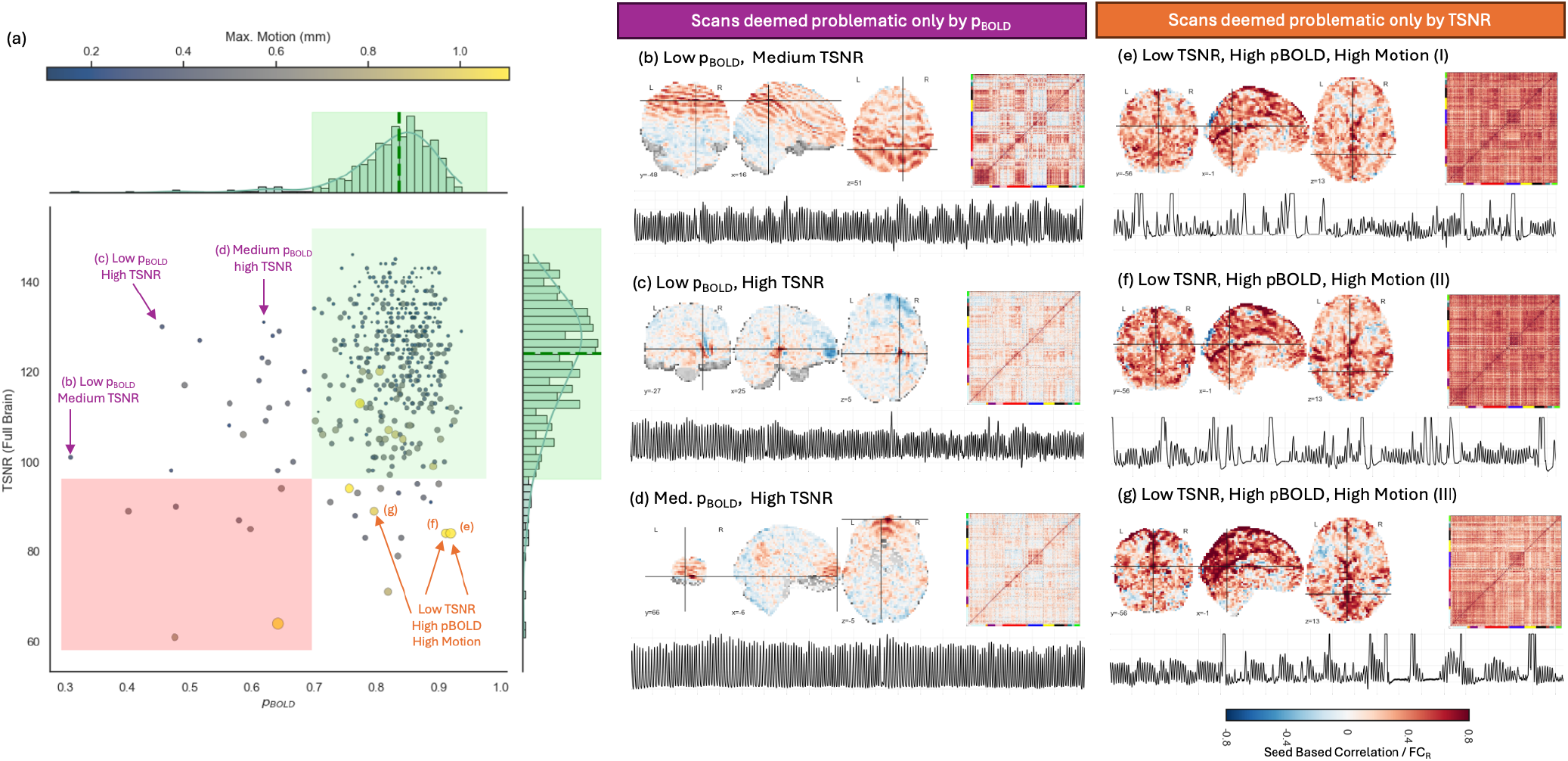
Identification of problematic scans with TSNR and p_BOLD_. (a) Scatter plot of TSNR vs. p_BOLD_ for the Basic pipeline and no NORDIC. Each point represents a scan with color and size conveying information about its maximum head motion. On the top and the right of the scatter we show the corresponding distributions of TSNR and p_BOLD_ for the whole dataset. The respective medians are marked with green dashed lines. The respective green shaded regions over the distribution correspond to the range [median – 2.5 * absolute mean difference, median + 2.5 * absolute mean difference]. The green shaded region of the scatter plot highlights the area occupied by scans deemed of good quality using both metrics. The red shaded region of the scatter plot highlights the area occupied by scans deemed of bad quality using both metrics. Arrows indicate scans to be explored in more detail. Next to each arrow is an indicator of the sub-panel associated with each of these scans. (b-d) Seed-based correlation (left), full brain FC matrix (right) and respiratory traces for scans deemed ok in terms of TSNR, but not in terms of p_BOLD_. for a scan with both low TSNR and p_BOLD_. (e-g) Same as (b-d) but for scans deemed ok in terms of p_BOLD_, but not in terms of TSNR.

We observe that the largest proportion of scans in the evaluation dataset are in the right-upper corner of the scatter plot (green shaded region), showcasing an overall high quality for this dataset. We also observe a few scans in the lower left quadrant (red shaded region) with both low *p_BOLD_* and TSNR. These quadrants represent scans where *p_BOLD_* and TSNR provide similar assessments of data quality.

We also observe scans outside these two quadrants. A subset of scans with high TSNR but low *p_BOLD_* sit on the upper right quadrant of the scatter plot. Figures 5b-d show a seed correlation map, a full brain *FC*_*R*_ matrix, and respiration trace for three scans in this region. Figure 5b is the scan with the lowest *p_BOLD_* (*p_BOLD_*=0.32 and TSNR=101). The seed correlation map reveals the presence of an artifact affecting large portions of the brain. The full brain *FC*_*R*_ matrix shows a clear network delineation, but also higher than usual levels of positive and negative inter-network correlation. Respiration is typical for this scan. Next, figures 5c-d show the same information for two scans with TSNR higher than the median TSNR of the whole sample, but low *p_BOLD_* values. In both cases, seed-based correlation analyses revealed the presence of artifacts, although these seem more spatially constrained than the one in Figure 5b. Full brain *FC*_*R*_ matrices show a clear network structure. Respiratory traces are quite regular. In summary, detailed exploration of representative scans likely to be deemed good in terms of TSNR, but problematic from a p_BOLD_ perspective, reveals that indeed these scans contain artifactual signals with the potential to negatively alter *FC*_*R*_ patterns.

Finally, figures 5e-g show equivalent visualizations of scans in the lower right quadrant of figure 5a. These are scans deemed problematic by TSNR, but not by p_BOLD_. For all three scans, seed-based maps with seed located in a high vascular volume region reveal the presence of widespread physiological contamination likely associated with prominent respiratory or cardiac variability. Full brain *FC*_*R*_ matrices show ubiquitous high positive correlation values consistent with an artifact of this nature. Respiratory traces confirm that subjects had quite irregular respiration patterns during these scans.

#### Properties of the Global Signal

Next, we evaluate two properties of the GS to understand why TSNR and *p_BOLD_* provide discrepant views regarding the always controversial GSR. The two ways we characterize the GS are: its likelihood of being BOLD via its *kappa* and *rho* statistic, and its variance explained by respiratory and cardiac-related nuisance regressors.

The *kappa* and *rho* statistics form the basis of TE-dependent analyses methods such as tedana (DuPre et al., 2021). Higher *kappa* indicates more BOLD contributions, while higher *rho* indicates more non-BOLD contributions to a time series. Figure 6a depicts the distributions of the GS’ *kappa* and *rho* statistics for all 439 scans. Of these, 381 scans (which account for 87% of scans) have higher *kappa* than *rho* – more BOLD than non-BOLD contributions (green shaded area of the scatter plot in Figure 6a). While *rho* statistics for the GS narrowly concentrate around its median value of 21.4 (red histogram to the right of Figure 6.a), *kappa* statistics are distributed over a much larger range of values (green histogram above in Figure 6a). In summary, in the majority of scans the GS have the characteristics of a BOLD signal.

**Figure 6.**
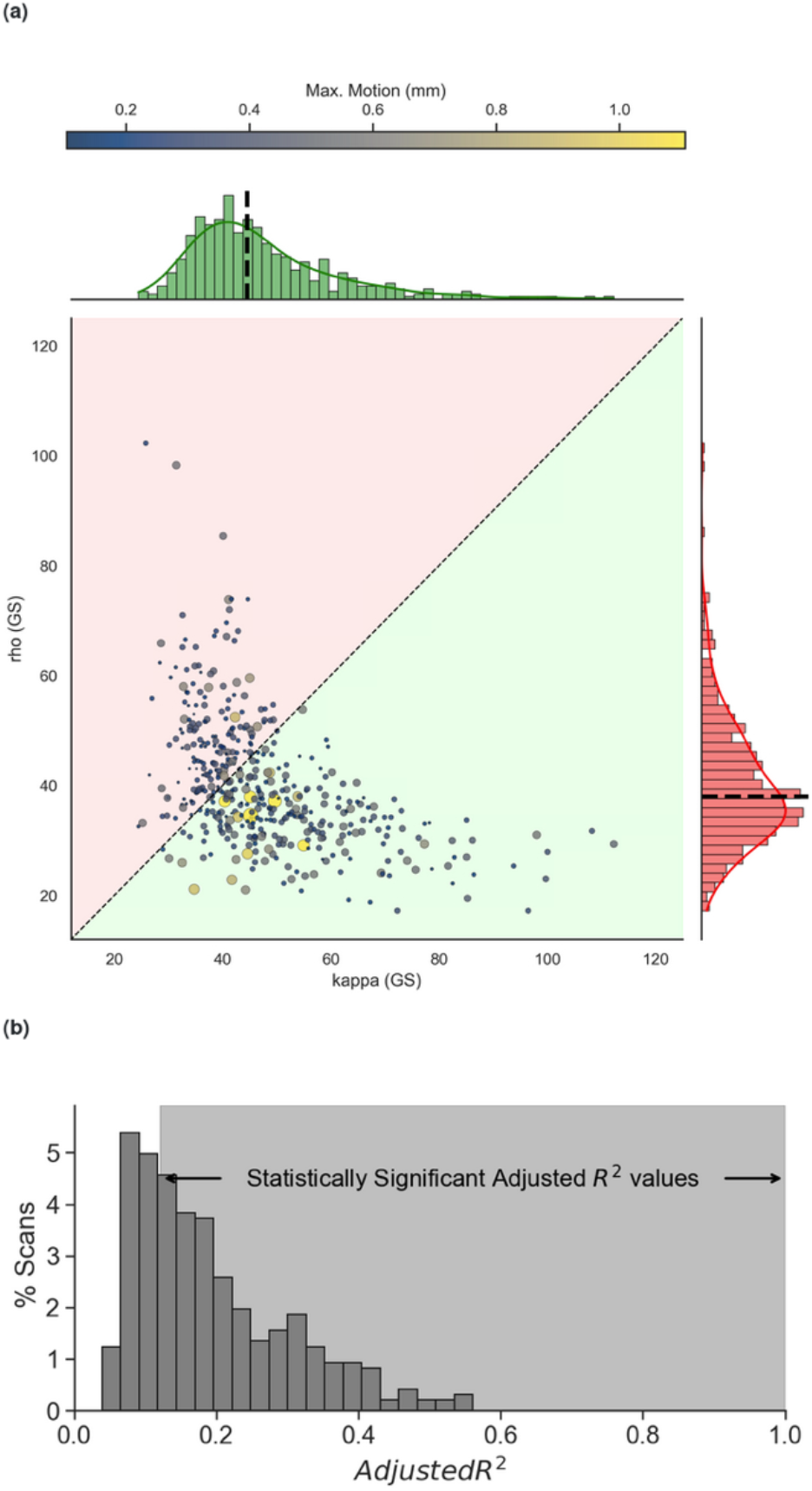
Kappa and Rho of the global signal for all scans. Each point represents a scan. Size and color of the scan represent the maximum head motion of the scan. Distributions of kappa and rho for the whole sample are shown above and to the right of the scatter plot, respectively. (b) Distribution of variance explained (adjusted R^2^) in the global signal by physiological regressors. Values that are statistically significant (p<0.05) are those in the light graded region.

Because not all BOLD fluctuations are of neural origin, and variability in respiratory and cardiac function can also induce BOLD fluctuations, it is important to understand the contribution of these two physiological processes to our observations. Figure 6b shows the distribution of explained variance (adjR^2^) in the GS by physiological regressors. Statistically significant values (p<0.05) relative to a null distribution with 10,000 permutations are those in the right shared region. 67% of scans fall in this region, yet of these, only 5 scans show adjR^2^>0.5. Overall, this suggests that the BOLD content of the GS cannot be primarily explained in terms of cardiac and respiratory functions.

#### Prediction of Fluid IQ scores

Figure 7 shows a summary of prediction of fluid IQ using CPM. Prediction accuracy was significantly lower for GSR than for Basic or Tedana denoising for both scanning sessions, independently of whether m-NORDIC was applied (Figures 7a-b). Prediction accuracy was significantly higher for tedana preprocessing compared to Basic pre-processing in all instances except for session 1 with m-NORDIC, where no significant difference was found (Figure 7a).

**Figure 7.**
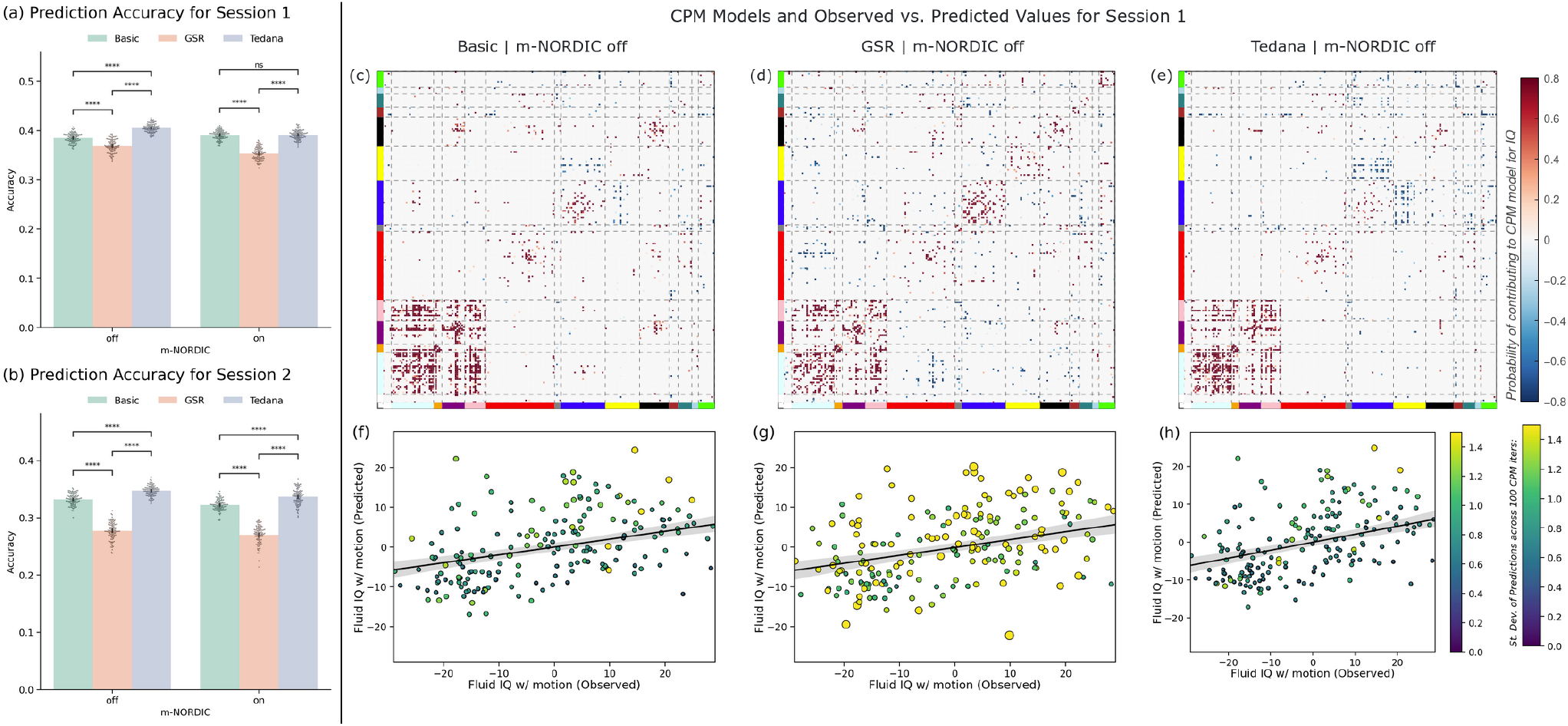
Connectome Predictive Modeling Results. (a) Prediction accuracy across for 100 attempts using the first scanning session for all pre-processing pipelines. Accuracy reported in terms of the Pearson’s correlation between observed and predicted IQ values after residualization with mean motion. Each dot represents a run of the CPM algorithm with a different initialization random seed. Bar height represents mean accuracy and error bars represent the 95% confidence interval. Statistical significance estimated using the Mann Whitney test (**** p<10^-5^, ns not significant). (b) Same as (a) but using the second scanning session as input features to the CPM algorithm. (c) CPM model for predicting fluid IQ using the first scanning session pre-processed with the Basic pipeline and no NORDIC. Edges found to positively correlate with fluid IQ are shown in red, those to correlate negatively in blue. The strength of the coloring indicates how often an edge was part of the CPM model across all 100 iterations. The solid line represents the linear fit to the data, and the shaded region the 95% confidence interval associated with this linear fit. (d) Same as (c) but for data pre-processed using the GSR pipeline and no NORDIC. (e) Same as (c) but for data pre-processed using the tedana pipeline and no NORDIC. (f) Scatter plot of average observed vs. predicted across all 100 iterations of the CPM algorithm using the first scanning session, with Basic pre-processing and no NORDIC. Each point represents a subject. The color and size of the points show the standard deviation of the predicted value across the 100 iterations. (g) Same as (f) for GSR and no NORDIC. (h) Same as (f) for tedana and no NORDIC.

Resulting CPM models, namely the set of edges that significantly correlate with fluid IQ, were quite similar across pre-processing pipelines (Figures 7c-e illustrate those for the three pipelines with no m-NORDIC). That said, for tedana, a larger number of edges are identified as having a negative correlation with fluid IQ (Figure 7e). Such edges correspond primarily to edges among regions of the Fronto-parietal Task Control network and the Visual network.

Finally, figures 7f-h depict the scatter plots of observed vs. predicted fluid IQ values for the three pre-processing pipelines that do not use m-NORDIC. In these plots, each dot represents a subject, and the color and size of the dot represents the standard deviation of predicted fluid IQ for that subject across the 100 CPM iterations. A higher variability in prediction values (i.e., lower precision) is clearly observed for the GSR pipeline (Figure 7g).

## Discussion

This work introduces and evaluates *p_BOLD_*, a new QA metric that leverages additional information that is available from ME-fMRI. Using two separate datasets we show: 1) that *p_BOLD_* accurately captures the likelihood that data are dominated by BOLD fluctuations; 2) how *p_BOLD_* complements TSNR when comparing pre-processing pipelines for seeking to identify problematic fMRI scans; and 3) that, as a QC measure, a higher *p_BOLD_* is a marker for better phenotypic prediction based on resting-state FC. Additionally, by relying on the observation that GSR enhances TSNR but lowers *p_BOLD_*, we provide some additional insights regarding the BOLD content of the GS and how its removal adversely affects data quality.

### TSNR and p_BOLD_ provide complementary information

TSNR is defined as the ratio of the mean signal over time divided by the standard deviation. Etched in this definition are two assumptions: 1) mean signal intensity is of interest to the experimenter, and 2) all fluctuations around the mean are uninformative and therefore considered noise. The first assumption always holds true when seeking to make inferences about neural function because BOLD fluctuations are proportional to average signal levels (see (Krüger and Glover, 2001b) for details). Unfortunately, the second assumption only applies when working with phantom objects (e.g., the fBIRN QA phantom) free of physiological fluctuations. Therefore, using phantom TSNR to evaluate acquisition protocols (Zwaag et al., 2012) or scanner stability (Kayvanrad et al., 2021) is straight forward: higher and stable TSNR is always better. Doing so for in-vivo data is more nuisance because, despite TSNR being indeed a valuable marker of data quality, TSNR can increase by unintended removal of fluctuations of interest (i.e., neuronally induced BOLD fluctuations). To avoid this unsought scenario, *p_BOLD_* purposely quantifies how much BOLD fluctuations dominate over other fluctuations in the data.

This divergence regarding how TSNR and *p_BOLD_* define “noise” explains the discrepancies observed here when evaluating pre-processing pipelines and, later on, when identifying problematic scans.

Regarding pipeline evaluation, figures 4.a illustrates that GSR produces TSNR increases when compared to Basic denoising. At the same time figure 4.b shows the opposite behavior in terms of *p_BOLD_*. TSNR improvement for GSR happens because including the GS as an additional nuisance regressor reduces the temporal standard deviation while leaving the mean signal untouched. Such an observation is not incompatible with increases in *p_BOLD_* should the GS be primarily BOLD in origin. Figure 6.a confirms that to be the case: 87% scans in the “evaluation” dataset have GS with higher *kappa* than *rho*; meaning their GS is primarily BOLD. In other words, in this sample, for most runs GSR removes BOLD-weighted signal.

Comparatively, figures 4.a,b also show that *tedana* produces data with higher TSNR and higher *p_BOLD_* relative to the other two pipelines. This demonstrates that a physics-informed selection of nuisance regressors—as *tedana* does—outperforms GSR in producing data with enhanced levels of BOLD fluctuations relative to all else. This does not automatically mean that *tedana* generates data free of confounds. Non-neural physiological processes (e.g., respiration and cardiac function) can contribute noise of BOLD nature to the data and *p_BOLD_* alone cannot detect such occurrences (see more on this topic below).

Next, figure 5a shows how TSNR and *p_BOLD_* can also provide complementary information when evaluating the quality of individual scans. For many scans TSNR and *p_BOLD_* provided congruent assessments, meaning both metrics suggested the presence or absence of issues. Yet, for approximately 10% of scans, we found that TSNR and *p_BOLD_* provided discrepant assessments (those sitting in uncolored regions in Figure 5a). First, we observed 21 scans occupying the upper right quadrant (high TSNR, low p_BOLD_). This suggests that although these scans have good temporal stability (as indicated by TSNR), the primary source of fluctuations in them corresponds to changes in net magnetization. Further inspection of several of these scans using AFNI’s *Instacorr* (Reynolds et al., 2023) confirmed this to be the case. Importantly, these observations exemplify how *p_BOLD_* can help identify problematic scans that would pass undetected using TSNR.

Additionally, 17 scans were found in the lower right quadrant of Figure 5a, which exhibit low TSNR but high *p_BOLD_*. Furthermore, several of these scans showed high motion estimates and their *FC*_*R*_ matrices were clearly contaminated by high correlation values everywhere. In a nutshell, *p_BOLD_* failed to identify these instances (3.8% of the sample) as problematic. The reason for this is that these scans were contaminated by respiratory-variability artifacts, which can be partially BOLD in nature. This was confirmed by seed-based correlation maps for the junction of the straight sinus and the vein of Galen—an area of high venous volume— that revealed a spatial pattern of correlations consistent with this type of physiological artifact: correlation with other highly vascularized regions such as the inferior occipital cortex, and the posterior cingular cortex (Birn et al., 2008, 2006). Irregular breathing patterns in the respiratory recordings further support this interpretation. Although this situation applied to a very small portion of the sample, it is key to notice it, because it highlights one limitation of *p_BOLD_*, and of TE-dependence analysis in general, which is that physiological processes that modulate the amount of CO_2_ in blood, which in turn alter blood flow (Greve et al., 2013; Wise et al., 2004), induce BOLD fluctuations with a TE-dependence profile indistinguishable from that of neuronally-induced BOLD fluctuations. Because of this, a sound interpretation of *p_BOLD_* requires having at hand information about cardiac and respiratory processes during the scans. Although not common practice in fMRI research, concurrent in-scanner respiratory and cardiac traces can be easily acquired using the pneumatic belt and plethysmograph readily available in most MRI scanners, or can be inferred in a data-driven manner using deep learning (Salas et al., 2021) and other analytical methods (Aslan et al., 2019).

### Global Signal Regression lowers *p_BOLD_* and phenotypic prediction power

Including the GS as a regressor improves TSNR but decreases *p_BOLD_* relative to Basic denoising. As already discussed, this is because the GS in this sample is primarily BOLD, as indicated by the distribution of *kappa* and *rho* statistics for the GS (Figure 6a). That said, is this BOLD predominance in the GS due to neural activity or other physiological processes (e.g., respiration variability)? To address this, we focused on a subset of scans from the evaluation sample for which concurrent physiological recordings (cardiac and respiratory) were available. We used these physiological traces and the *RETROICOR* (Glover et al., 2000) and *RVT* (Birn et al., 2006) models to generate nuisance regressors that model the most common effects of cardiac, respiration and respiration variability in fMRI recordings. We explored how well all these physiological regressors explained the GS. We found that significant amounts of variance in the GS could be explained by physiological regressors in 67% of the sample; yet the average explained variance was 20% (in line with prior reports of 20% (Power et al., 2017) and 31% (Liu et al., 2017)) and only above 50% for 5 scans. This suggests that the BOLD fluctuations in the GS are a mixture of neural and non-neural physiological sources, and that GSR will inevitably remove the contributions of neural processes of interest. Yet this observation must be interpreted with care given that RETROICOR+RVT regressors (such as the ones used here) are known to have limited explanatory power. For example, Hocke et al. (2016) have estimated that optimized physiological regressors derived from peripheral NIRs recordings can explain up to 16% more variance that the RETROICOR+RVT regressors explored here. That said, even when considering this potential gap in explanatory power, physiological regressors would still only partially account for a limited amount of variance in most of our sample. As such, its regression will still remove the contribution of neural processes of potential interest. Prior literature has cautioned against using the GS as a regressor because it artifactually induces negative correlations (Murphy et al., 2009), can alter group differences (Saad et al., 2012), it correlates strongly with spontaneous fluctuations in local field potentials (Schölvinck et al., 2010), and its amplitude also correlates with EEG measures of vigilance (Wong et al., 2013) (please see (Liu et al., 2017) for an in-depth discussion of sources of information in the GS).

Our results further support the views that GSR can be problematic as it decreases the ratio of BOLD fluctuations relative to other sources, and this is achieved at the expense of removing BOLD contributions which cannot be fully ascribed to cardiac and/or respiratory function.

Further support for this view that GSR can change data and its connectivity distributions in undesired ways comes from the observation that GSR produced worse performance (in terms of accuracy and precision) when predicting fluid IQ. Fluid IQ quantifies the ability to “solve problems, think and act quickly… and plays an important role in adapting to novel situations in everyday life” (Akshoomoff et al., 2013). As such, fluid IQ is more likely linked to each subject’s underlying neural activity and connectivity patterns than to their respiratory and cardiac patterns while lying still in the scanner bore. Therefore, the decrease in predicting fluid IQ following GSR suggests that regressing the GS results in the removal and alteration of FC patterns that meaningfully encode the complex set of cognitive abilities that contribute to this phenotypic marker. That said, it is important to note that these results are at odds with conclusions from two recent studies looking at how GSR and other denoising methods affect brain-wide associations (Li et al., 2019; Pavlovich et al., 2025). While both studies report that GSR improves brain-wide associations, in the case of Pavlovich et al. they report that observed improvements are marginal. Because our study and theirs differ in many aspects, including the figure of merit used, the baseline condition, the brain parcellation, and acquisition parameters; it is difficult to pinpoint the reason/s leading to the opposing conclusions. That said, importantly for our discussion, *p_BOLD_* changes across pre-processing pipelines more directly match changes in prediction accuracy than TSNR does. This suggests that, independently of why GSR reduced prediction performance here and not in the other studies, *p_BOLD_* was able to more accurately capture that outcome than TSNR alone.

### Tedana and multi-echo denoising

One additional observation is that tedana denoising also outperformed Basic and GSR pipelines in terms of prediction accuracy and precision (Figure 7). This suggests that a more targeted approach to the generation of nuisance regressors, in the case of tedana by characterizing their TE-dependence, can improve the strength of brain-wise associations without incurring the interpretative challenges associated with GSR. This agrees with prior work showing improvements in trial-level associations between BOLD activity levels and response time in an object identification task (Gotts et al., 2024). Our results also extend an already long list of benefits associated with ME-fMRI, and tedana denoising in particular, such as increased statistical power for task-based fMRI (Lombardo et al., 2016), reduced data needs for precision functional mapping (Lynch et al., 2020), improved sensitivity for the detection of neural activity of unknown timing (Caballero-Gaudes et al., 2019), improved removal of artifacts associated with overt speech paradigms (Gilmore et al., 2022) and motor-task paradigms (Reddy et al., 2024).

### Inclusion of NORDIC and interactions with pre-processing pipelines

Despite using magnitude-only NORDIC denoising—because phase data was not available— m-NORDIC significantly lowered thermal noise in both datasets. This thermal noise reduction translated into significant increases in TSNR and *p_BOLD_* across all pre-processing pipelines (Figure 4a-b, Table 2 and Suppl. Figure 3). Of note, improvements in TSNR were larger in magnitude than those observed with *p_BOLD_*. This is likely because of how *p_BOLD_* is defined, which focuses exclusively on how empirical across-echo *FC*_*C*_ matrices behave in relation to two theoretical extreme signal regimes (BOLD and So dominant sources). *p_BOLD_* is not intended to quantify any relationship with thermal noise content. In fact, our theoretical derivation for *p_BOLD_* assumes that thermal noise is negligible relative to BOLD and S_o_ sources. Estimation of thermal noise pre- and post-m-NORDIC are in the range of 3 to 6, which are substantially lower levels of variability than those observed inside the brain and therefore suggests that our assumption holds for these data (Suppl. Figure 2).

Running m-NORDIC prior to any other pre-processing step resulted in mixed results regarding ability to predict fluid IQ. For session 1 scans, m-NORDIC significantly improved prediction accuracy for Basic denoising but worsened it for GSR and tedana pipelines. For session 2 scans, prediction accuracy worsened for all three pipelines. In summary, in 5 out of 6 scenarios, m-NORDIC resulted in a small, though significant, reduction in prediction accuracy. Although NORDIC has been reported to increase the fMRI sensitivity to small neural effects (Faes et al., 2024; Moeller et al., 2021), specially at high spatial resolutions where thermal noise dominates, it has been noted that NORDIC can mistakenly remove signal of interest (Faes et al., 2024). We believe this is the case here. Examination of the mean, standard deviation and correlation patterns of the differential timeseries (without m-NORDIC minus with m-NORDIC) in the two scans from session 1 that resulted in the largest percentage of thermal noise being removed reveals two quite distinct situations (Suppl. Figure 4). For one scan, although the mean difference map contains structure, the standard deviation map and correlation map to the visual cortex do not. Conversely, for the other scan clear spatial structure can be observed in all maps. This suggests that while m-NORDIC might have removed thermal noise in all scans, for a subset of them, it has also removed other signal components. This could be because we used m-NORDIC, as opposed to phase-based NORDIC, or because we used the default NORDIC hyper-parameters, instead of optimizing them for our specific data. Prior work evaluating NORDIC for volume-sensitive vascular occupancy (VASO) data (Huber et al., 2017) has reported that NORDIC performance and robustness against eliminating signals of interest can vary depending on hyperparameter selection (Knudsen et al., 2025). Our results suggest the same is true for BOLD acquisitions. Importantly for our discussion regarding *p_BOLD_*, our results suggest that *p_BOLD_* might be more sensitive to this issue than TSNR, given that *p_BOLD_* better reflect the behavior observed for m-NORDIC in the prediction task.

### Limitations and Future work

This work presents *p_BOLD_*, a new QA metric specifically targeted for ME-fMRI, and showcases a few applications for it: pre-processing pipeline evaluation, identification of problematic scans, forecast accuracy for brain-wide associations. While our results prove *p_BOLD_* is effective at these tasks, *p_BOLD_* is not free of interpretational pitfalls, especially when it comes to contamination by non-neural BOLD processes. Fortunately, these BOLD contaminating processes have somehow predictable spatiotemporal profiles (i.e., overlap with high vascular volume regions) that can exploited to improve *p*_*BOLD*._ The use of spatial heuristics has proven successful for the automatic removal of motion artifacts (Pruim et al., 2015). Future work should evaluate if such heuristics could be used to create an additional complementary QA metric to be used with *p_BOLD_* and TSNR.

Here, we illustrate how *p_BOLD_* can be used to assess the quality of a dataset preprocessed in six different ways. In principle, *p_BOLD_* could also support the selection of scan-specific preprocessing pipelines that maximize BOLD content. For example, *p_BOLD_* could be incorporated into loss functions that iteratively select nuisance regressors and preprocessing steps to minimize non-BOLD fluctuations relative to those of BOLD origin. Future work should investigate how to define such loss functions so that maximizing BOLD content does not come at the expense of other important considerations, such as excessive loss of degrees of freedom or the production of data dominated by physiological BOLD effects that may not accurately reflect localized neural activity

One procedural caveat with *p_BOLD_’s* current formulation is that requires a brain parcellation. In this work we selected the Power 264 brain parcellation (Power et al., 2011) because of its wide use in the neuroimaging literature; but in principle, *p_BOLD_* can be calculated using other whole-brain parcellations. Future work should carefully evaluate *p_BOLD_’s* dependence on parcellation selection. We expect some level of dependence will be observed given that parcellations can substantially differ in terms of spatial coverage (e.g., full grey matter ribbon vs. small spheres at particular locations). Based on this expectation, we recommend that those looking to incorporate *p_BOLD_* into their QA protocols choose the same parcellation they plan to use in their main analyses. By doing so, *p_BOLD_* observations will be optimally tailored to their study needs and objectives.

*p_BOLD_* was defined using the assumption that thermal noise is negligible in comparison to BOLD and net magnetization effects (see theory section). This assumption is certain in most scenarios where ME acquisitions are used (e.g., 3T data with voxels of dimensions of 2×2×2mm or larger), yet it might not hold for higher spatial resolutions (please see (Bodurka et al., 2007) for additional details regarding the relationship between voxel size and thermal noise). Future work should evaluate how *p_BOLD_* behaves in such high-resolution regimes given the steady historical evolution of fMRI towards higher spatial and temporal resolutions.

Similarly, we also use a mono-exponential decay signal model for our theoretical derivations (Eq. 1). This is a common assumption when working with a limited number of echo times (Kundu et al., 2014; Peltier and Noll, 2002), as it is the case here. However, it is worth noting that more complex models have been proposed. For example, a biexponential model (Speck et al., 2001), and several multi-compartment models (Hoogenraad et al., 2001) have been proposed to better capture the multi-tissue composition of typical fMRI voxels. Future work should test if alternative formulations of p_BOLD_ based on these advanced signal models could provide more accurate estimates of data quality.

The theory underlying *p_BOLD_* focuses on how inter-regional covariance behaves across echo times, but the same logic also applies to intra-regional signal variance. Therefore, it should be possible to define alternative formulations of *p_BOLD_* that do not require to estimate full brain *FC*_*C*_ matrices. Such alternative formulations could be defined at the ROI level—which would still require a brain parcellation—or directly at the voxel-level. A voxel-level formulation, in addition to not requiring a parcellation, could provide users with QA observations at two different levels of detail. An average *p_BOLD_* across voxels might be a good indicator of overall scan quality. Additionally, a spatial map of voxel-wise *p_BOLD_* might help identify locations where data issues concentrate. Having such maps can be quite beneficial, as TSNR maps have proven before, because one can more precisely tailor scan selection based on whether problematic regions overlap with those targeted by a specific scientific inquiry. Developing this form of objective-driven QA tools—known as contextual QA (Teves et al., 2023; Wang and Strong, 1996)—is key for fMRI given the cost associated with data acquisition.

## Conclusions

The benefits of ME-fMRI acquisitions for clinical and basic neuroscience are becoming widely recognized by the functional neuroimaging community and adoption of this form of acquisitions is growing fast in recent years. ME-fMRI data are not immune to fMRI artifacts and its quality must be monitored, as is the case with single-echo data. One key advantage of ME-fMRI is that by using TE-dependence analysis one can distinguish between BOLD and S_o_ fluctuations present in the data. Prior work has demonstrated that this property of ME-fMRI data can be exploited to automatically identify noise sources, improve sensitivity to true underlying effects and the quality of deconvolution methods. Here, we further increase this list of benefits by showing how echo time dependence analysis can also be used to define a new QA metric specific to ME-fMRI that informs us explicitly about the desirable dominance of BOLD fluctuations above all else. We demonstrate that because *p_BOLD_* can better differentiate between noise sources, it can outperform more traditional metrics (e.g., TSNR) at tasks such as pre-processing evaluation and detection of problematic scans. In that way, *p_BOLD_* provides a meaningful additional tool that we recommend adding to current QA protocols for ME-fMRI data. Now, because not all BOLD fluctuations are necessarily neural, we discuss at length the interpretational limitations that *p_BOLD_* has when it comes to detecting datasets contaminated by strong wide-spread BOLD fluctuations of physiological origin (e.g., due to stereotypical variability in cardiac and respiratory function). We demonstrate how its combined use with concurrent physiological recordings can help ameliorate this interpretational issue. Finally, we discuss possible enhancements to the *p_BOLD_* metric.

## Data and Code availability

Two datasets are included in this work. The discovery dataset (N=7) can be shared upon request. Neuroimaging data for the evaluation dataset (N=439) is a publicly available through OpenNeuro: https://openneuro.org/datasets/ds003592. Demographic and behavioral data for the evaluation dataset are available within the Open Science Framework project “Goal-Directed Cognition in Older and Younger Adults” (http://osf.io/yhzxe/).

Code used in these analyses is available in a public github repository: https://github.com/nimh-sfim/me_staticfc. This repository includes a stand-alone python program (*compute_pBOLD.py*) that can estimate *p_BOLD_* for a multi-echo dataset given ROI timeseries per echo, and a list of echo times.

## Author Contributions

J.G.C.: Conceptualization, Data Curation, Formal Analysis, Investigation, Methodology, Software, Visualization, Writing – Original Draft; C.C.G.: Conceptualization, Methodology, Writing—Review & Editing; D.A.H.: Conceptualization, Methodology, Writing—Review & Editing; P.A.B.: Funding acquisition, Resources, Supervision, Writing—Review & Editing.

## Acknowledgements

This research was made possible by the support of the National Institute of Mental Health Intramural Research Program (ZIAMH002783). C.C.G. contributions to this work were supported by the Project PID2023-149410b-100 funded by the Spanish Ministry of Science, Innovation and Universities (MCIU /AEI/10.13039/501100011033/FEDER.UE), the BCBL “Severo Ochoa” excellence accreditation (CEX2020-001010/AEI/10.13039/501100011033) and the Basque Government (BERC 2022–2025). The study was completed in compliance with the National Institutes of Health Clinical Center protocol ID 93-M-0170 (ClinicalTrials.gov ID: NCT00001360). This work utilized the computational resources of the NIH HPC Biowulf cluster (https://hpc.nih.gov).

This research was supported by the Intramural Research Program of the National Institutes of Health (NIH). The contributions of the NIH authors were made as part of their official duties as NIH federal employees, are in compliance with agency policy requirements, and are considered Works of the United States Government. However, the findings and conclusions presented in this paper are those of the authors and do not necessarily reflect the views of the NIH or the U.S. Department of Health and Human Services.

## Supplementary Figures

**Supplementary Figure 1.**
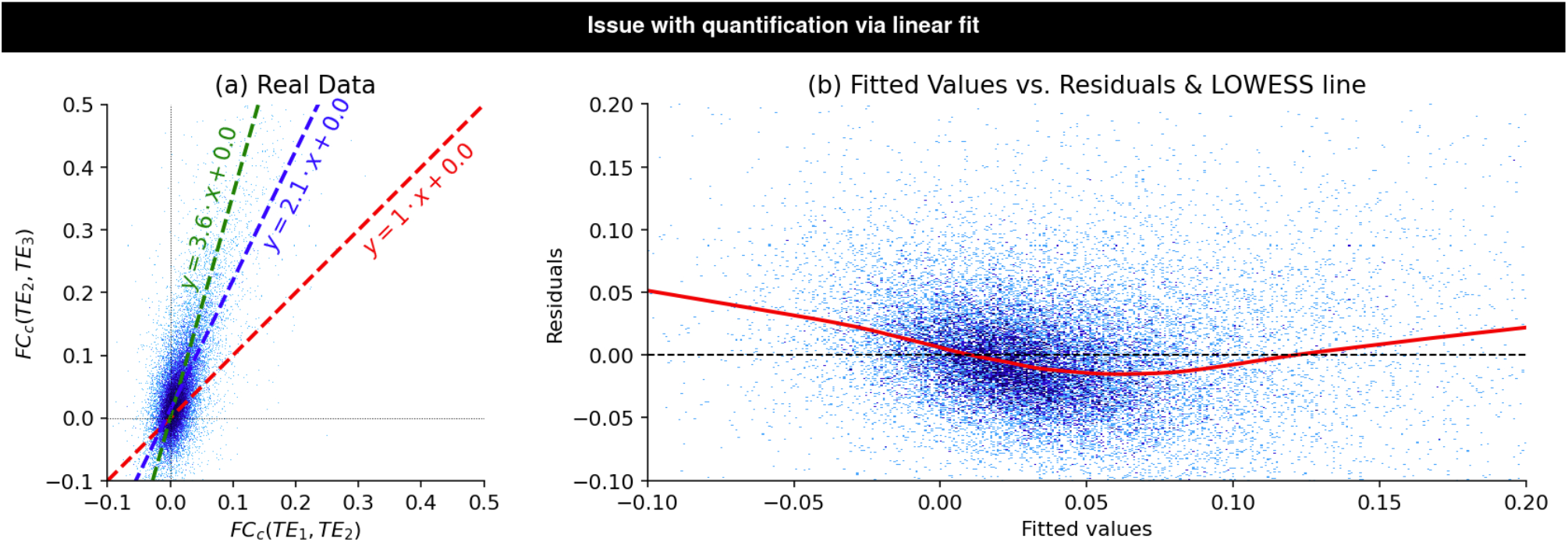
Representative dataset showing why a simple linear fit to the data is not a robust way to quantify where a given *FC*_*C*_matrix sits relative to the two signal regimes of interest. (a) Scatter plot of *FC*_*C*_ for two separate pairs of echo times. Edges are depicted as blue dots. Green dashed line shows expected behavior for BOLD dominated data. Red dashed line (identity line) shows expected behavior for non-BOLD dominated data. Blue dashed line shows the linear fit to the point cloud associated with input *FC*_*C*_. Although the data primarily sits over the green dashed line, the linear fit might suggest differently. (b) Scatter plot of residual values for the linear fit as a function of fitted values. We can observe that residuals are not homogenously distributed, which signals heteroscedasticity. The red line corresponds to the locally weighted scatter plot smoothing (LOWESS) fit to the cloud point.

**Supplementary Figure 2.**
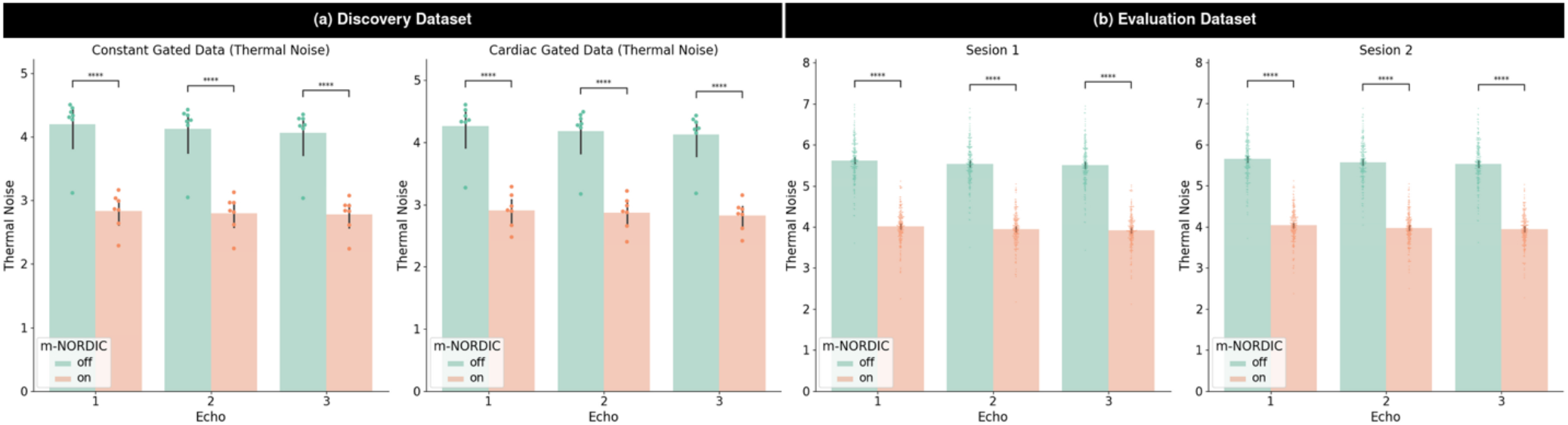
Reduction in thermal noise obtained using m-NORDIC. (a) Results for the discovery dataset with constant-gated scans on the left and cardiac-gated scans on the right. (b) Results for the evaluation dataset with scans from session 1 on the left and scans from session 2 on the right. In all plots colored bars indicate average values across the sample, error bars the 95% confidence interval and dots the results for each individual scan. Statistical annotations correspond to a double-sided paired T-test (****: p < 1e^-4^).

**Supplementary Figure 3.**
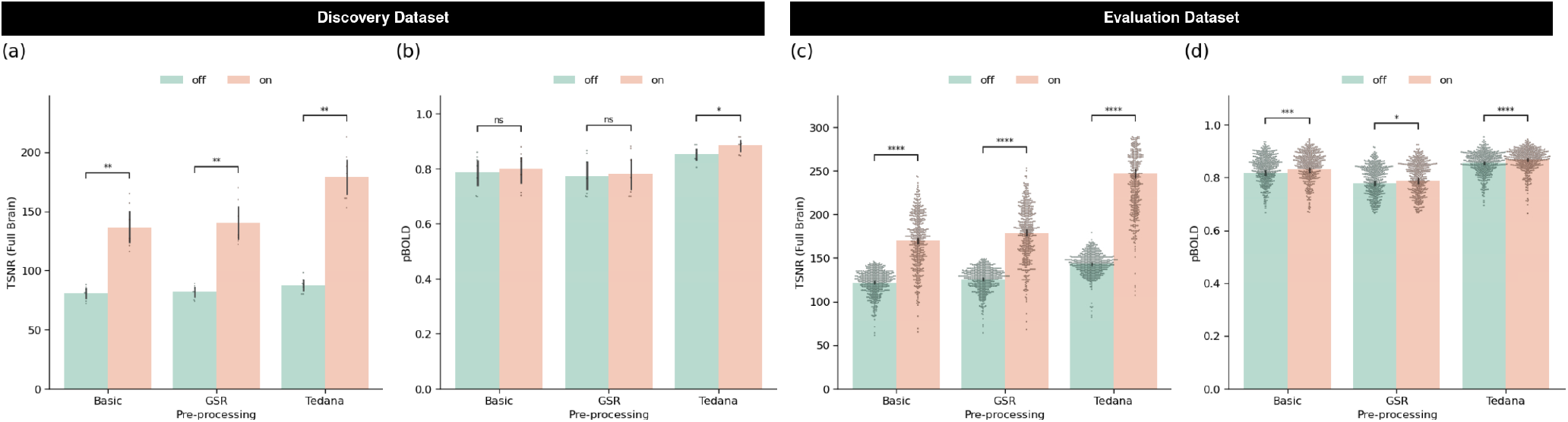
Changes in TSNR and *p_BOLD_* when applying m-NORDIC. (a) and (b) TSNR and *p_BOLD_* results for the discovery dataset, respectively. It includes only constant-gated data. (c) and (d) TSNR and *p_BOLD_* results for the evaluation dataset. It includes data from both sessions. Statistical annotations correspond to a double-sided Mann-Whitney Test (ns = not significant; * p<0.05; ** p<0.01; *** p<0.001; **** p< 1e^-4^.

**Supplementary Figure 4.**
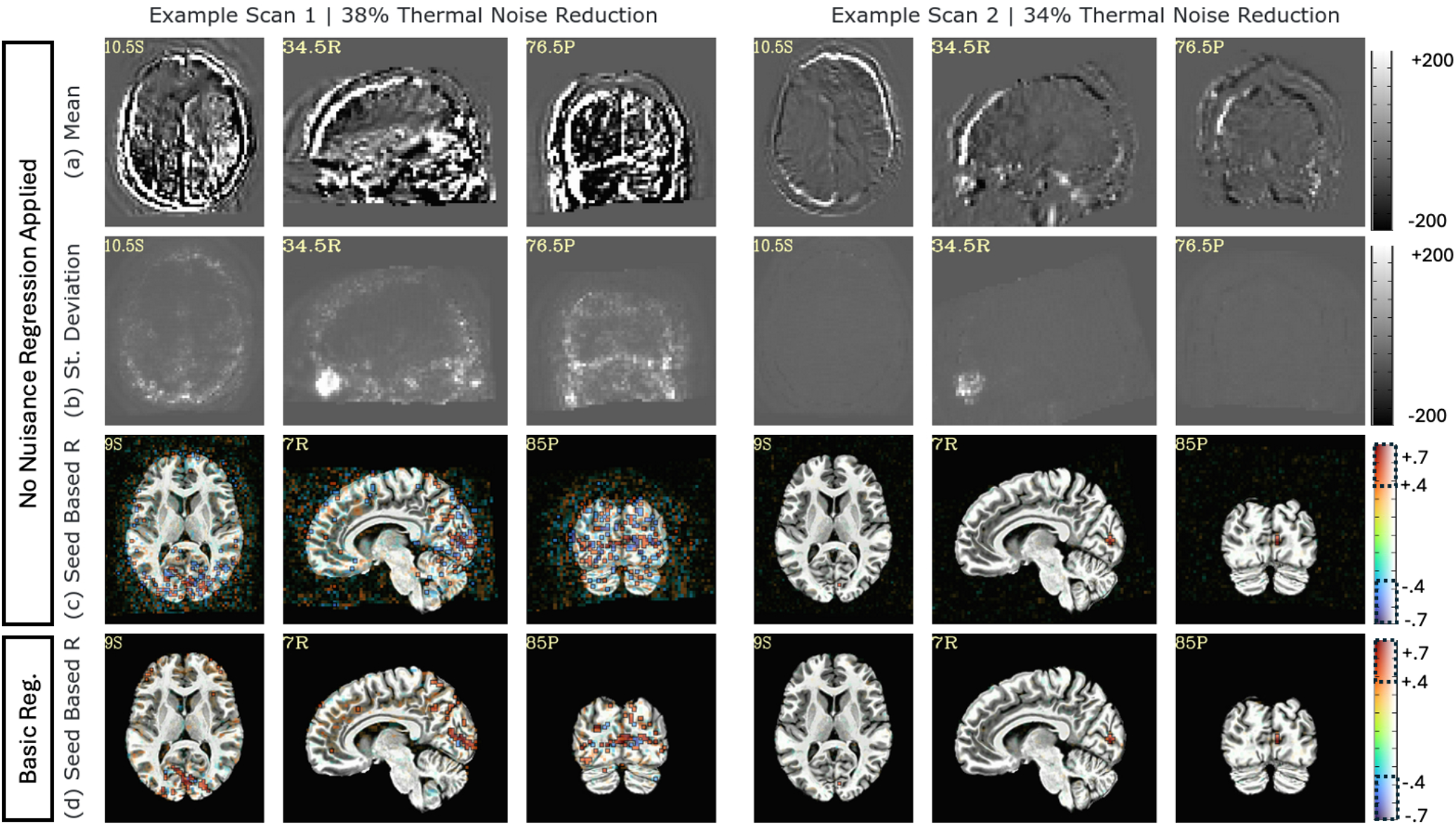
Effects of m-NORDIC denoising for the two scans that had the largest amount of thermal noise removed. All maps in this figure were computed using the differential timeseries between having applied or not applied NORDIC. Results are presented following spatial normalization into MNI space (top three columns) and following Basic Nuisance regression (bottom column). In theory, if m-NORDIC only removed random thermal noise, no spatial or correlational structure should be observed. (a) Mean across time of the differential timeseries for two exemplary scans. Both maps show clear structure, although the magnitude of that structure is much higher on the right. (b) Standard deviation across time of the differential timeseries. The map on the right shows a clear spatial structure, while for the scan on the left, structure is only appreciated near the eyes. (c) Seed-based correlation for a seed voxel in visual cortex prior to any nuisance regression. Clear correlational structure is observed on the right scan, but not the left. (d) Seed-based correlation for a seed voxel in visual cortex following Basic nuisance regression. Clear correlational structure is observed on the right scan, but not the left.

## References

Akshoomoff, N., Beaumont, J.L., Bauer, P.J., Dikmen, S.S., Gershon, R.C., Mungas, D., Slotkin, J., Tulsky, D., Weintraub, S., Zelazo, P.D., Heaton, R.K., 2013. VIII. NIH TOOLBOX COGNITION BATTERY (CB): COMPOSITE SCORES OF CRYSTALLIZED, FLUID, AND OVERALL COGNITION. Monogr. Soc. Res. Child Dev. 78, 119–132. 10.1111/mono.12038

Aslan, S., Hocke, L., Schwarz, N., Frederick, B., 2019. Extraction of the cardiac waveform from simultaneous multislice fMRI data using slice sorted averaging and a deep learning reconstruction filter. Neuroimage 198, 303–316. 10.1016/j.neuroimage.2019.05.049

Bandettini, P.A., Jesmanowicz, A., Wong, E.C., Hyde, J.S., 1993. Processing strategies for time-course data sets in functional mri of the human brain. Magn. Reson. Med. 30, 161–173. 10.1002/mrm.1910300204

Behzadi, Y., Restom, K., Liau, J., Liu, T.T., 2007. A component based noise correction method (CompCor) for BOLD and perfusion based fMRI. Neuroimage 37, 90–101. 10.1016/j.neuroimage.2007.04.042

Birn, R., Smith, M., Jones, T., Bandettini, P., 2008. The respiration response function: the temporal dynamics of fMRI signal fluctuations related to changes in respiration 40.

Birn, R.M., Diamond, J.B., Smith, M.A., Bandettini, P.A., 2006. Separating respiratory-variation-related fluctuations from neuronal-activity-related fluctuations in fMRI. NeuroImage 31, 1536–48. 10.1016/j.neuroimage.2006.02.048

Bodurka, J., Ye, F., Petridou, N., Murphy, K., Bandettini, P.A., 2007. Mapping the MRI voxel volume in which thermal noise matches physiological noise—Implications for fMRI. NeuroImage 34, 542–549. 10.1016/j.neuroimage.2006.09.039

Breusch, T.S., Pagan, A.R., 1979. A Simple Test for Heteroscedasticity and Random Coefficient Variation. Econometrica 47, 1287. 10.2307/1911963

Caballero-Gaudes, C., Moia, S., Panwar, P., Bandettini, P.A., Gonzalez-Castillo, J., 2019. A deconvolution algorithm for multi-echo functional MRI: Multi-echo Sparse Paradigm Free Mapping. Neuroimage 116081. 10.1016/j.neuroimage.2019.116081

Chang, L.-C., Rohde, G.K., Pierpaoli, C., 2005. An automatic method for estimating noise-induced signal variance in magnitude-reconstructed magnetic resonance images. Méd. Imaging 2005: Image Process. 1136–1142. 10.1117/12.596008

Cohen, A.D., Chang, C., Wang, Y., 2021. Using multiband multi-echo imaging to improve the robustness and repeatability of co-activation pattern analysis for dynamic functional connectivity. NeuroImage 243, 118555. 10.1016/j.neuroimage.2021.118555

Cox, R.W., Jesmanowicz, A., 1999. Real-time 3D image registration for functional MRI. Magn. Reson. Med. 42, 1014–1018. 10.1002/(sici)1522-2594(199912)42:6<1014::aid-mrm4>3.0.co;2-f

Dubois, J., Galdi, P., Paul, L.K., Adolphs, R., 2018. A distributed brain network predicts general intelligence from resting-state human neuroimaging data. Philosophical Transactions of the Royal Society B 373. 10.1098/rstb.2017.0284

DuPre, E., Salo, T., Ahmed, Z., Bandettini, P., Bottenhorn, K., Caballero-Gaudes, C., Dowdle, L., Gonzalez-Castillo, J., Heunis, S., Kundu, P., Laird, A., Markello, R., Markiewicz, C., Moia, S., Staden, I., Teves, J., Uruñuela, E., Vaziri-Pashkam, M., Whitaker, K., Handwerker, D., 2021. TE-dependent analysis of multi-echo fMRI with tedana. J. Open Source Softw. 6, 3669. 10.21105/joss.03669

Evans, J.W., Kundu, P., Horovitz, S.G., Bandettini, P.A., 2015. Separating slow BOLD from non-BOLD baseline drifts using multi-echo fMRI. NeuroImage 105, 189–197. 10.1016/j.neuroimage.2014.10.051

Faes, L.K., Lage-Castellanos, A., Valente, G., Yu, Z., Cloos, M.A., Vizioli, L., Moeller, S., Yacoub, E., Martino, F.D., 2024. Evaluating the effect of denoising submillimeter auditory fMRI data with NORDIC. Imaging Neurosci. 2, imag–2–00270. 10.1162/imag_a_00270

Finn, E., Shen, X., Scheinost, D., Rosenberg, M., 2014. Functional connectome fingerprinting: identifying individuals using patterns of brain connectivity. Nature Neuroscience 18. 10.1038/nn.4135

Fischl, B., 2012. FreeSurfer. Neuroimage 62, 774–81. 10.1016/j.neuroimage.2012.01.021

Frederick, B. deB., Nickerson, L.D., Tong, Y., 2012. Physiological denoising of BOLD fMRI data using Regressor Interpolation at Progressive Time Delays (RIPTiDe) processing of concurrent fMRI and near-infrared spectroscopy (NIRS). Neuroimage 60, 1913–1923. 10.1016/j.neuroimage.2012.01.140

Friedman, L., Glover, G.H., 2006. Report on a multicenter fMRI quality assurance protocol. J. Magn. Reson. Imaging 23, 827–839. 10.1002/jmri.20583

Gati, J.S., Menon, R.S., Uğurbil, K., Rutt, B.K., 1997. Experimental determination of the BOLD field strength dependence in vessels and tissue. Magnetic Resonance in Medicine 38, 296–302. 10.1002/mrm.1910380220

Gilmore, A.W., Agron, A.M., González-Araya, E.I., Gotts, S.J., Martin, A., 2022. A Comparison of Single- and Multi-Echo Processing of Functional MRI Data During Overt Autobiographical Recall. Front. Neurosci. 16, 854387. 10.3389/fnins.2022.854387

Glen, D.R., Taylor, P.A., Buchsbaum, B.R., Cox, R.W., Reynolds, R.C., 2020. Beware (Surprisingly Common) Left-Right Flips in Your MRI Data: An Efficient and Robust Method to Check MRI Dataset Consistency Using AFNI. Front. Neuroinformatics 14, 18. 10.3389/fninf.2020.00018

Glover, G., Li, T., Ress, D., 2000. Image-based method for retrospective correction of physiological motion effects in fMRI: RETROICOR 44. 10.1002/1522-2594(200007)44:1<162::aid-mrm23>3.0.co;2-e

Gonzalez-Castillo, J., Hoy, C.W., Handwerker, D.A., Roopchansingh, V., Inati, S.J., Saad, Z.S., Cox, R.W., Bandettini, P.A., 2015. Task Dependence, Tissue Specificity, and Spatial Distribution of Widespread Activations in Large Single-Subject Functional MRI Datasets at 7T. Cerebral Cortex 25, 4667–77. 10.1093/cercor/bhu148

Gonzalez-Castillo, J., Panwar, P., Buchanan, L.C., Caballero-Gaudes, C., Handwerker, D.A., Jangraw, D.C., Zachariou, V., Inati, S., Roopchansingh, V., Derbyshire, J.A., Bandettini, P.A., 2016. Evaluation of multi-echo ICA denoising for task based fMRI studies: Block designs, rapid event-related designs, and cardiac-gated fMRI. NeuroImage 141. 10.1016/j.neuroimage.2016.07.049

Gonzalez-Castillo, J., Saad, Z.S., Handwerker, D.A., Inati, S.J., Brenowitz, N., Bandettini, P.A., 2012. Whole-brain, time-locked activation with simple tasks revealed using massive averaging and model-free analysis. Proceedings of the National Academy of Sciences 109. 10.1073/pnas.1121049109

Gotts, S.J., Gilmore, A.W., Martin, A., 2024. Harnessing slow event-related fMRI to investigate trial-level brain-behavior relationships during object identification. Front. Hum. Neurosci. 18, 1506661. 10.3389/fnhum.2024.1506661

Greve, D.N., Brown, G.G., Mueller, B.A., Glover, G., Liu, T.T., Network, F.B.R., 2013. A Survey of the Sources of Noise in fMRI. Psychometrika 78, 396–416. 10.1007/s11336-012-9294-0

Hocke, L.M., Tong, Y., Lindsey, K.P., Frederick, B. de B., 2016. Comparison of peripheral near-infrared spectroscopy low-frequency oscillations to other denoising methods in resting state functional MRI with ultrahigh temporal resolution. Magnetic Resonance in Medicine 76, 1697–1707. 10.1002/mrm.26038

Hoogenraad, F.G.C., Pouwels, P.J.W., Hofman, M.B.M., Reichenbach, J.R., Sprenger, M., Haacke, E.M., 2001. Quantitative differentiation between BOLD models in fMRI. Magn. Reson. Med. 45, 233–246. 10.1002/1522-2594(200102)45:2<233::aid-mrm1032>3.0.co;2-w

Huber, L., Handwerker, D.A., Jangraw, D.C., Chen, G., Hall, A., Stüber, C., Gonzalez-Castillo, J., Ivanov, D., Marrett, S., Guidi, M., Goense, J., Poser, B.A., Bandettini, P.A., 2017. High-Resolution CBV-fMRI Allows Mapping of Laminar Activity and Connectivity of Cortical Input and Output in Human M1. Neuron 96, 1253-1263.e7. 10.1016/j.neuron.2017.11.005

Jo, H., Saad, Z.S., Simmons, K.W., Milbury, L.A., Cox, R.W., 2010. Mapping sources of correlation in resting state FMRI, with artifact detection and removal. NeuroImage 52, 571–82. 10.1016/j.neuroimage.2010.04.246

Kayvanrad, A., Arnott, S.R., Churchill, N., Hassel, S., Chemparathy, A., Dong, F., Zamyadi, M., Gee, T., Bartha, R., Black, S.E., Lawrence-Dewar, J.M., Scott, C.J.M., Symons, S., Davis, A.D., Hall, G.B., Harris, J., Lobaugh, N.J., MacQueen, G., Woo, C., Strother, S., Investigators, the O.F., Investigators, the C.-B., 2021. nnResting state fMRI scanner instabilities revealed by longitud inal phantom scans in a multi-center study. NeuroImage 237, 118197. 10.1016/j.neuroimage.2021.118197

Knudsen, L., Vizioli, L., Martino, F.D., Faes, L.K., Handwerker, D.A., Moeller, S., Bandettini, P.A., Huber, L., 2025. NORDIC denoising on VASO data. Front. Neurosci. 18, 1499762. 10.3389/fnins.2024.1499762

Korponay, C., Janes, A.C., Frederick, B.B., 2024. Brain-wide functional connectivity artifactually inflates throughout functional magnetic resonance imaging scans. Nat. Hum. Behav. 8, 1568–1580. 10.1038/s41562-024-01908-6

Krüger, G., Glover, G.H., 2001a. Physiological noise in oxygenation-sensitive magnetic resonance imaging: Physiological Noise in MRI. Magnet Reson Med 46, 631–637. 10.1002/mrm.1240

Krüger, G., Glover, G.H., 2001b. Physiological noise in oxygenation-sensitive magnetic resonance imaging. Magn. Reson. Med. 46, 631–637. 10.1002/mrm.1240

Kundu, P., Inati, S.J., Evans, J.W., Luh, W.-M., Bandettini, P.A., 2011. Differentiating BOLD and non-BOLD signals in fMRI time series using multi-echo EPI. Neuroimage 60, 1759–70. 10.1016/j.neuroimage.2011.12.028

Kundu, P., Santin, M.D., Bandettini, P.A., Bullmore, E.T., Petiet, A., 2014. Differentiating BOLD and non-BOLD signals in fMRI time series from anesthetized rats using multi-echo EPI at 11.7T. NeuroImage 102, 861–874. 10.1016/j.neuroimage.2014.07.025

Li, J., Kong, R., Liégeois, R., Orban, C., Tan, Y., Sun, N., Holmes, A.J., Sabuncu, M.R., Ge, T., Yeo, B.T.T., 2019. Global signal regression strengthens association between resting-state functional connectivity and behavior. NeuroImage 196, 126–141. 10.1016/j.neuroimage.2019.04.016

Liu, T.T., Nalci, A., Falahpour, M., 2017. The global signal in fMRI: Nuisance or Information? NeuroImage 150, 213–229. 10.1016/j.neuroimage.2017.02.036

Lombardo, M.V., Auyeung, B., Holt, R.J., Waldman, J., Ruigrok, A.N.V., Mooney, N., Bullmore, E.T., Baron-Cohen, S., Kundu, P., 2016. Improving effect size estimation and statistical power with multi-echo fMRI and its impact on understanding the neural systems supporting mentalizing. NeuroImage 142, 55–66. 10.1016/j.neuroimage.2016.07.022

Lynch, C.J., Elbau, I., Liston, C., 2021. Improving precision functional mapping routines with multi-echo fMRI. Curr Opin Behav Sci 40, 113–119. 10.1016/j.cobeha.2021.03.017

Lynch, C.J., Power, J.D., Scult, M.A., Dubin, M., Gunning, F.M., Liston, C., 2020. Rapid Precision Functional Mapping of Individuals Using Multi-Echo fMRI. Cell Reports 33, 108540. 10.1016/j.celrep.2020.108540

Makowski, C., Lepage, M., Evans, A.C., 2019. Head motion: the dirty little secret of neuroimaging in psychiatry. J. Psychiatry Neurosci. 44, 62–68. 10.1503/jpn.180022

Menon, R.S., Ogawa, S., Tank, D.W., Uğurbil, K., 1993. Tesla gradient recalled echo characteristics of photic stimulation-induced signal changes in the human primary visual cortex. Magnetic resonance in medicine 30, 380–6.

Moeller, S., Pisharady, P.K., Ramanna, S., Lenglet, C., Wu, X., Dowdle, L., Yacoub, E., Uğurbil, K., Akçakaya, M., 2021. NOise reduction with DIstribution Corrected (NORDIC) PCA in dMRI with complex-valued parameter-free locally low-rank processing. NeuroImage 226, 117539. 10.1016/j.neuroimage.2020.117539

Moia, S., Termenon, M., Uruñuela, E., Chen, G., Stickland, R.C., Bright, M.G., Caballero-Gaudes, C., 2021. ICA-based denoising strategies in breath-hold induced cerebrovascular reactivity mapping with multi echo BOLD fMRI. NeuroImage 233, 117914. 10.1016/j.neuroimage.2021.117914

Morfini, F., Whitfield-Gabrieli, S., Nieto-Castañón, A., 2023. Functional connectivity MRI quality control procedures in CONN. Front. Neurosci. 17, 1092125. 10.3389/fnins.2023.1092125

Murphy, K., Birn, R.M., Handwerker, D.A., Jones, T.B., Bandettini, P.A., 2009. The impact of global signal regression on resting state correlations: Are anti-correlated networks introduced? NeuroImage 44, 893–905. 10.1016/j.neuroimage.2008.09.036

Oakes, T.R., Johnstone, T., Walsh, K.S.O., Greischar, L.L., Alexander, A.L., Fox, A.S., Davidson, R.J., 2005. Comparison of fMRI motion correction software tools. NeuroImage 28, 529–543. 10.1016/j.neuroimage.2005.05.058

Parrish, T.B., Gitelman, D.R., LaBar, K.S., Mesulam, M. -Marsel, 2000. Impact of signal-to-noise on functional MRI. Magn. Reson. Med. 44, 925–932. 10.1002/1522-2594(200012)44:6<925::aid-mrm14>3.0.co;2-m

Pavlovich, K., Pang, J., Holmes, A., Constable, T., Fornito, A., 2025. The efficacy of resting-state fMRI denoising pipelines for motion correction and behavioural prediction. Imaging Neurosci. 3, IMAG.a.97. 10.1162/imag.a.97

Peltier, S.J., Noll, D.C., 2002. T2* Dependence of Low Frequency Functional Connectivity. Neuroimage 16, 985–992. 10.1006/nimg.2002.1141

Power, J.D., Barnes, K.A., Snyder, A.Z., Schlaggar, B.L., Petersen, S.E., 2012. Spurious but systematic correlations in functional connectivity MRI networks arise from subject motion. Neuroimage 59, 2142–2154.

Power, J.D., Cohen, A.L., Nelson, S.M., Wig, G.S., Barnes, K.A., Church, J.A., Vogel, A.C., Laumann, T.O., Miezin, F.M., Schlaggar, B.L., Petersen, S.E., 2011. Functional Network Organization of the Human Brain. Neuron 72, 665–678. 10.1016/j.neuron.2011.09.006

Power, J.D., Plitt, M., Laumann, T.O., Martin, A., 2017. Sources and implications of whole-brain fMRI signals in humans. Neuroimage 146, 609–625. 10.1016/j.neuroimage.2016.09.038

Provins, C., MacNicol, E., Seeley, S.H., Hagmann, P., Esteban, O., 2023. Quality control in functional MRI studies with MRIQC and fMRIPrep. Front. Neuroimaging 1, 1073734. 10.3389/fnimg.2022.1073734

Pruim, R.H.R., Mennes, M., Rooij, D. van, Llera, A., Buitelaar, J.K., Beckmann, C.F., 2015. ICA-AROMA: A robust ICA-based strategy for removing motion artifacts from fMRI data. NeuroImage 112, 267–277. 10.1016/j.neuroimage.2015.02.064

Reddy, N.A., Zvolanek, K.M., Moia, S., Caballero-Gaudes, C., Bright, M.G., 2024. Denoising task-correlated head motion from motor-task fMRI data with multi-echo ICA. Imaging Neurosci. 2, 1–30. 10.1162/imag_a_00057

Reeder, S.B., Atalar, E., Bolster, B.D., McVeigh, E.R., 1997. Quantification and reduction of ghosting artifacts in interleaved echo-planar imaging. Magn. Reson. Med. 38, 429–439. 10.1002/mrm.1910380312

Reynolds, R.C., Glen, D.R., Chen, G., Saad, Z.S., Cox, R.W., Taylor, P.A., 2024. Processing, evaluating and understanding FMRI data with afni_proc.py. arXiv. 10.48550/arxiv.2406.05248

Reynolds, R.C., Taylor, P.A., Glen, D.R., 2023. Quality control practices in FMRI analysis: Philosophy, methods and examples using AFNI. Front. Neurosci. 16, 1073800. 10.3389/fnins.2022.1073800

Saad, Z.S., Gotts, S.J., Murphy, K., Chen, G., Jo, H.J., Martin, A., Cox, R.W., 2012. Trouble at Rest: How Correlation Patterns and Group Differences Become Distorted After Global Signal Regression. Brain Connect. 2, 25–32. 10.1089/brain.2012.0080

Salas, J.A., Bayrak, R.G., Huo, Y., Chang, C., 2021. Reconstruction of respiratory variation signals from fMRI data. Neuroimage 225, 117459. 10.1016/j.neuroimage.2020.117459

Schölvinck, M.L., Maier, A., Ye, F.Q., Duyn, J.H., Leopold, D.A., 2010. Neural basis of global resting-state fMRI activity. Proceedings of the National Academy of Sciences 107. 10.1073/pnas.0913110107

Shen, X., Finn, E.S., Scheinost, D., Rosenberg, M.D., Chun, M.M., Papademetris, X., Constable, R.T., 2017. Using connectome-based predictive modeling to predict individual behavior from brain connectivity. Nature Protocols 12. 10.1038/nprot.2016.178

Speck, O., Ernst, T., Chang, L., 2001. Biexponential modeling of multigradient-echo MRI data of the brain. Magn. Reson. Med. 45, 1116–1121. 10.1002/mrm.1147

Spreng, R.N., Setton, R., Alter, U., Cassidy, B.N., Darboh, B., DuPre, E., Kantarovich, K., Lockrow, A.W., Mwilambwe-Tshilobo, L., Luh, W.-M., Kundu, P., Turner, G.R., 2022. Neurocognitive aging data release with behavioral, structural and multi-echo functional MRI measures. Sci. Data 9, 119. 10.1038/s41597-022-01231-7

Teves, J.B., Gonzalez-Castillo, J., Holness, M., Spurney, M., Bandettini, P.A., Handwerker, D.A., 2023. The art and science of using quality control to understand and improve fMRI data. Front Neurosci-switz 17, 1100544. 10.3389/fnins.2023.1100544

Triantafyllou, C., Polimeni, J.R., Wald, L.L., 2011. Physiological noise and signal-to-noise ratio in fMRI with multi-channel array coils. NeuroImage 55, 597–606. 10.1016/j.neuroimage.2010.11.084

Uruñuela, E., Gonzalez-Castillo, J., Zheng, C., Bandettini, P., Caballero-Gaudes, C., 2024. Whole-brain multivariate hemodynamic deconvolution for functional MRI with stability selection. Méd. Image Anal. 91, 103010. 10.1016/j.media.2023.103010

Wang, R.Y., Strong, D.M., 1996. Beyond Accuracy: What Data Quality Means to Data Consumers. J. Manag. Inf. Syst. 12, 5–33. 10.1080/07421222.1996.11518099

Wilcox, R.R., Barbey, A.K., 2023. Connectome-based predictive modeling of fluid intelligence: evidence for a global system of functionally integrated brain networks. Cereb. Cortex 33, 10322–10331. 10.1093/cercor/bhad284

Wise, R.G., Ide, K., Poulin, M.J., Tracey, I., 2004. Resting fluctuations in arterial carbon dioxide induce significant low frequency variations in BOLD signal. NeuroImage 21, 1652–1664. 10.1016/j.neuroimage.2003.11.025

Wong, C.W., Olafsson, V., Tal, O., Liu, T.T., 2013. The amplitude of the resting-state fMRI global signal is related to EEG vigilance measures. NeuroImage 83. 10.1016/j.neuroimage.2013.07.057

Zwaag, W. van der, Marques, J.P., Kober, T., Glover, G., Gruetter, R., Krueger, G., 2012. Temporal SNR characteristics in segmented 3D-EPI at 7T. Magn. Reson. Med. 67, 344–352. 10.1002/mrm.23007

